# Dynamic acetylation of a conserved lysine impacts glycerol kinase activity and abundance in the haloarchaeon *Haloferax volcanii*

**DOI:** 10.1101/2025.07.03.662939

**Authors:** Karol M. Sanchez, Manasa Addagarla, Heather Judd, Xin Wang, Julie Maupin-Furlow

## Abstract

*Haloferax volcanii* is a halophilic archaeon that preferentially utilizes glycerol as a carbon source, placing glycerol kinase (GK, *glpK*) at the center of its metabolism. In contrast to bacterial GKs, which are often regulated by allosteric inhibition, *H. volcanii* GK lacks this mode of control, indicating alternative regulatory mechanisms. Here, we show that lysine acetylation of *H. volcanii* GK enhances its activity and abundance during growth on glycerol, with K153 identified as the primary site of modification. Structural modeling and comparative genomics revealed that K153 resides in a conserved flexible loop common to haloarchaeal GKs. Carbon shifts from glucose to glycerol led to increased activity and enrichment of the K153-acetylated form, as determined by AQUA-MS. GK and the acetylation mimic K153Q supported growth on glycerol, while the non-acetylatable K153R variant did not. Thermal shift analysis showed that the K153R substitution reduced GK stability, while K153Q had no effect. Size exclusion chromatography indicated that GK is predominantly dimeric but forms a tetramer when purified from glycerol-grown cells and assayed with glycerol - coinciding with the highest K153 acetylation levels. Kinetic analysis revealed that K153 acetylation is required to maintain cooperative substrate binding, with the non-acetylatable K153R variant exhibiting a loss of allosteric behavior. The GNAT-family acetyltransferase Pat2 was found to acetylate GK at K153, and *Δpat2* mutants exhibited reduced GK protein abundance, linking Pat2 to regulation of GK. These results identify a dynamic, carbon source-responsive lysine acetylation mechanism that modulates GK, highlighting lysine acetylation as a key component of haloarchaeal metabolic regulation.

**IMPORTANCE:** Post-translational modifications allow microorganisms to rapidly adapt their metabolism to changing environmental conditions. Here, we uncover a carbon source-dependent acetylation mechanism that regulates GK activity and abundance in the halophilic archaeon *H. volcanii*. Unlike bacterial systems, where allosteric inhibitors control GK, *H. volcanii* relies on lysine acetylation to fine-tune and enhance enzymatic function based on nutrient availability. Our findings highlight acetylation at a conserved lysine as a key modulator of archaeal carbon metabolism, linking environmental signals directly to enzymatic activity and cellular fitness. This work expands our understanding of extremophile metabolic regulation and reveals how archaea deploy unique strategies to survive and thrive in environments that shift in carbon availability.

## INTRODUCTION

Glycerol metabolism plays a critical role in the survival and environmental adaptation of haloarchaea in hypersaline environments (1). In *Haloferax volcanii*, glycerol is the preferred carbon source over glucose, with glycerol kinase (GK) serving as the key enzyme for glycerol utilization (2). Our recent work characterized the biochemical properties of the *H. volcanii* GK enzyme, demonstrating its robust catalytic activity under extreme conditions, including exposure to organic solvents, elevated temperatures, and high salinity (3). However, the regulatory mechanisms governing GK activity in *H. volcanii* remained unknown.

In bacteria such as *Escherichia coli*, GK is regulated allosterically by fructose-1,6-bisphosphate (FBP) and the EIIA component of the phosphotransferase system (PTS) (4–6). In contrast, archaeal GKs, including *H. volcanii* GK, lack the equivalent regulatory domains found in bacteria, suggesting that alternative mechanisms have evolved in archaea to modulate GK activity in response to environmental cues.

Upon exposure to glycerol, glucose catabolism in *H. volcanii* is repressed at both transcriptional and protein levels, indicating a carbon catabolite repression mechanism adapted to favor glycerol metabolism (7). This prioritization is enforced at the transcriptional level by GlpR, a DeoR-family repressor, which downregulates the expression of genes involved in glucose and fructose catabolism in the presence of glycerol (7). In contrast, the transcriptional regulator GfcR acts as an activator of sugar catabolism when glycerol is limited, promoting the expression of genes encoding enzymes involved in glucose and fructose degradation, including gluconate dehydratase, glyceraldehyde-3-phosphate dehydrogenase, and pyruvate kinase (8).

Beyond transcriptional regulation, recent omics approaches have expanded the understanding of post-translational modifications (PTMs) in *H. volcanii*, including lysine acetylation. A high-throughput mass spectrometry (MS) study of glycerol-grown *H. volcanii* has identified GK to be acetylated at K153, suggesting a potential role for lysine acetylation in the regulation of carbon metabolism (9). Acetylation at the analogous lysine residue was also reported in the GK of *Haloferax mediterranei*, implying conservation across halophilic archaea (10).

Acetylation can influence protein conformation, stability, solubility, interactions, and enzymatic activity by altering surface charge and hydrophobicity (11). Lysine acetylation can function as a reversible regulatory mechanism, capable of either activating or repressing enzyme activity depending on the context (12). In eukaryotes, acetylation is best known for activating histones (13), while in bacteria and archaea, its roles are more diverse and only recently being elucidated. Lysine acetylation does appear evolutionarily conserved and dynamically modulated in response to nutrient availability, acting as a metabolic rheostat (14, 15).

Despite increasing recognition of lysine acetylation as a key regulator of metabolism, the functional impact of this modification on haloarchaeal GK enzymes remains unclear. Here, we show that the abundance, activity, and thermal stability of *H. volcanii* GK are regulated by Pat2-mediated acetylation at K153 in a carbon source-dependent manner. Acetylation levels at this site correlate with GK enzymatic activity and support growth on glycerol. Structural modeling places K153 within a flexible loop conserved among most haloarchaeal GKs, suggesting a lineage-specific regulatory feature. These findings reveal a novel post-translational mechanism controlling archaeal metabolism, with broader implications for biotechnology and evolutionary biology.

## MATERIALS AND METHODS

### Strains and media

Strains, plasmids, and oligonucleotides used in this study are listed in **Table S1-S2**. *Escherichia coli* Top10 (Invitrogen) was used for routine cloning, and *E. coli* GM2163 (New England Biolabs) was used for propagation of unmethylated plasmids for *H. volcanii* transformations. *E. coli* strains were grown at 37 °C in Luria-Bertani (LB) medium supplemented with 100 μg/mL ampicillin and/or 34 µg/mL chloramphenicol as needed. *H. volcanii* strains were cultured at 42 °C in ATCC 974 complex medium (ATCC) or minimal media supplemented with glycerol (GlyMM), glucose (GluMM), or fructose (FruMM) at 20 mM each carbon source final concentration. Growth media were supplemented as needed with uracil (50 μg/mL), novobiocin (Nov, 0.1 μg/mL), 5-fluoroorotic acid (5-FOA, 50 μg/mL dissolved in DMSO), and isopropyl-β-D-1-thiogalactopyranoside (IPTG, 0.4 mM). For growth assays, colonies from solid media were inoculated into 5 mL of medium. Cells were subcultured at an initial optical density monitored at 600 nm (OD₆₀₀) of 0.02. Cell growth was monitored by an increase in OD₆₀₀, where 1 OD₆₀₀ unit equals 1× 10^9^ colony forming units (CFU)·ml^-1^ for all strains used in this study.

### Construction of *H. volcanii* strains

Markerless gene deletions were generated using the *pyrE2*-based “pop-in/pop-out” method as previously described (16). Flanking regions (∼500 bp) were amplified and cloned into the BamHI and HindIII sites of pTA131. The resulting plasmids were transformed into *H. volcanii* strains that included the *ΔpyrE2* mutation, and transformants were selected on uracil-deficient media (HV-CA+). Counter-selection was performed on 5-FOA-containing plates (ATCC). Mutants were identified by PCR screening and confirmed by Sanger sequencing using primers outside of the 500 bp flanking regions. *H. volcanii* H1207 (Δ*pyrE2* Δ*pitA)* was the parental strain used to generate the double mutant strain H1207 Δ*glpK* Δ*larC* (designated KM03). KM03 was constructed to eliminate co-purification of the endogenous His-rich LarC (HVO_2381, putative nickel insertion protein) **(Fig. S1)**. Additional strains derived from KM03, including Δ*pat1* (KM05), Δ*pat2* (KM06), Δ*sir2* (KM07), and Δ*elp3* (KM08), were generated to investigate the effects of acetyltransferase and deacetylase mutations on GK regulation *in vivo*.

### Plasmid construction

The plasmids used in this study are listed in **Table S2**. High-fidelity PCR was used to amplify the *H. volcanii glpK* gene (HVO_1541) using Phusion DNA polymerase (New England Biolabs) and cloned into two different expression systems. For native expression in *H. volcanii*, the *glpK* gene was inserted into the pJAM503 backbone modified for N-terminal His-tag fusion, generating plasmid pJAM4351. For recombinant expression in *E. coli* Rosetta (DE3), the gene was cloned into the pET15b vector, generating plasmid pJAM4360. Amplified inserts and vectors were digested with NdeI and BlpI restriction enzymes, purified using the Monarch PCR and DNA Cleanup Kit (New England Biolabs), ligated with T4 DNA ligase, and transformed into *E. coli* Top10 cells. Positive clones were screened by colony PCR and verified by Sanger DNA sequencing (Eurofins Genomics). Site-directed mutagenesis (SDM) to generate the K153R and K153Q variants of GK was performed by the SSPER and rrPCR methods previously described (17), followed by DpnI digestion and transformation into *E. coli* GM2163 for plasmid propagation. Verified plasmids were subsequently transformed into *H. volcanii* H1207 for native expression or *E. coli* Rosetta (DE3) for recombinant expression.

### Protein modeling and multiple sequence alignment

The predicted three-dimensional structure of *H. volcanii* GK (UniProt ID: D4GYI5, HVO_1541) was generated using AlphaFold2 (18) via the ColabFold implementation (19). Structural visualization and analysis were conducted using UCSF ChimeraX v1.5 (20). Multiple sequence alignment of 39 GK homologs from different genera from the *Halobacteria* class (**Dataset S1**), was performed using Clustal Omega (21), and results were visualized using ESPript 3.0 (22).

### GK expression and purification

His-tagged GK expression and purification from *H. volcanii* were carried out as previously described (3). Briefly, *H. volcanii* strains were grown to stationary phase, harvested, and lysed in buffer containing 50 mM HEPES (pH 7.5), 2 M NaCl, 5 mM β-mercaptoethanol, 40 mM imidazole, DNase I, and protease inhibitors. Lysates were clarified by centrifugation and incubated with Ni-NTA resin (Millipore Sigma) at 4 °C. Proteins were eluted with 250 mM imidazole in HEPES buffer and analyzed by SDS-PAGE. For *E. coli*-expressed recombinant His-GK (HvGK_Ec_), *E. coli* Rosetta (DE3) cells carrying pJAM4360 were induced with 0.4 mM IPTG at 25 °C overnight. HvGK_Ec_ was purified similarly, with buffer exchange to 2 M NaCl using Zeba spin desalting columns to enhance protein stability (23).

### Mapping acetylation sites and protein identification by LC-MS/MS analysis

The following strategy was used for mapping acetylation sites and identifying unknown proteins. Protein samples were purified by Ni-NTA affinity chromatography, separated by SDS-PAGE, stained with BioSafe Coomassie Brilliant Blue (CB), and excised from the gel prior to LC-MS/MS analysis. Samples included *H. volcanii* GK expressed from a plasmid with an N-terminal His-tag (His-GK) and purified from *H. volcanii* H1207-pJAM4351 cells grown in glycerol (GlyMM). An additional protein that co-purified from glucose-(GluMM-) grown *H. volcanii* H1207 cells carrying the empty vector (pJAM202c) was also analyzed. For comparison, recombinant His-GK was analyzed following heterologous expression from plasmid pJAM4360 in *E. coli* Rosetta (DE3) cells, grown in LB medium supplemented with chloramphenicol.

Excised protein bands were subjected to in-gel digestion using sequencing-grade trypsin or chymotrypsin (Promega, Madison, WI), following the manufacturer’s protocol with minor modifications. Gel bands were trimmed closely to reduce background and washed twice with nanopure water, followed by two 10-min washes with 1:1 (v/v) methanol: 50 mM ammonium bicarbonate. Dehydration was performed with 1:1 (v/v) acetonitrile: 50 mM ammonium bicarbonate. Gel slices were reduced with 25 mM dithiothreitol (DTT) in 100 mM ammonium bicarbonate for 30 min and alkylated with 55 mM iodoacetamide for 30 min in the dark. Following another round of washing and dehydration, protease digestion was initiated by rehydrating gels with 12 ng/µL trypsin in 0.01 % ProteaseMAX surfactant, followed by overlaying with 40 µL of 0.01 % ProteaseMAX in 50 mM ammonium bicarbonate. Digestion was allowed to proceed for 1 h with gentle shaking. Reactions were quenched with 0.5 % trifluoroacetic acid (TFA), and samples were either analyzed immediately or stored at –80 °C.

Peptides were analyzed by nanoLC-MS/MS using a Thermo Scientific Q Exactive HF Orbitrap mass spectrometer with an EASY-Spray nanosource and an UltiMate 3000 RSLCnano system (Thermo Scientific). Peptides were first loaded onto a PharmaFluidics mPAC C18 trapping column (10 mm) with 1 % buffer B (acetonitrile + 0.1 % formic acid) and desalted at a flow rate of 10 µL/min. Peptides were then eluted onto a 50 cm mPAC C18 analytical column and separated using a gradient from 1 % to 20 % buffer B over 60 min, then up to 45 % over 10 min (total run time 90 min). The flow rate was initially set at 750 nL/min for 15 min and reduced to 300 nL/min thereafter. Column temperature was maintained at 40 °C.

MS was operated in data-dependent acquisition mode using the Top15 method. Full scans (m/z 375–1575) were acquired at 30,000 resolutions, with AGC target of 3e6 and maximum injection time of 50 ms. The most intense ions were selected for MS/MS at 15,000 resolutions with a 2e5 AGC target and 55 ms maximum injection time. HCD fragmentation was performed using stepped normalized collision energy (NCE) of 28, with a 4.0 m/z isolation window. Singly charged ions and known isotopes were excluded. Dynamic exclusion was enabled (repeat count: 1; exclusion duration: 15 s). The siloxane ion at m/z 445.12003 served as an internal calibrant. A HeLa digest standard was analyzed regularly to ensure instrument performance, with a minimum acceptance threshold of 2700 protein IDs.

Raw MS/MS spectra were processed using Proteome Discoverer 3.2 (Thermo Fisher Scientific). Spectra were searched using the Sequest HT algorithm against *H. volcanii* Uniprot ID: UP000008243 supplemented with *E. coli* and common contaminant sequences. Search parameters included trypsin specificity, 10 ppm precursor ion tolerance, and 0.02 Da fragment ion tolerance. Carbamidomethylation of cysteine was set as a fixed modification; methionine oxidation, N-terminal acetylation, methionine loss (± acetylation), and lysine acetylation were included as variable modifications. Peptides were filtered for a 1 % false discovery rate using Percolator. Acetylated lysine residues were manually validated, and MS2 fragment spectra were inspected to confirm site localization.

### Protein quantification and SDS-PAGE analysis

Protein concentrations were determined by Bradford assay using bovine serum albumin (Bio-Rad Laboratories) as a standard. Samples were resolved by 8 or 12 % SDS–PAGE as indicated, and protein fractions were prepared for electrophoresis by mixing equal volume of Laemmli SDS sample buffer: 100 mM Tris-HCl (pH 6.8), 10 % (v/v) β-mercaptoethanol, 2 % (w/v) SDS, 10 % (v/v) glycerol, and 0.6 mg/mL bromophenol blue. The samples were boiled for 10 min, chilled on ice for 5 min, and centrifuged at 13,000 × *g* for 10 min. Precision Plus Protein Kaleidoscope molecular mass marker (Bio-Rad Laboratories) was the standard. The gels were stained with CB and imaged using an iBright Imaging System (Invitrogen) according to the manufacturer’s protocol.

### Immunoblot analysis

After SDS-PAGE, proteins were transferred to PVDF membranes (0.2 µm) (Amersham, Little Chalfont, UK) at 25 V for 7 min using the Trans-Blot Turbo Transfer System (Bio-Rad Laboratories) and Towbin buffer (as recommended by supplier). After protein transfer, the membranes were incubated with gentle rocking overnight at 4 °C. For His-tagged proteins, membranes were blocked in buffer composed of 5 % (w/v) skim milk powder in TBST [50 mM Tris-HCl, pH 7.6, 150 mM NaCl, and 0.1 % (v/v) Tween 20], and subsequently probed for 1 h at RT with HRP-conjugated 6×His-tag mouse monoclonal antibody (Proteintech Group, Inc, Rosemont, IL, U.S. Cat. No. HRP-660005) diluted 1:10,000 in blocking buffer. For lysine acetylation proteins, membranes were blocked in buffer composed of 5 % (w/v) BSA in TBST and subsequently probed for 1 h at RT with anti-acetyllysine rabbit mAb (PTM Bio, #PTM-105RM) as the primary antibody at 1:5,000 dilution and mouse anti-rabbit IgG-HRP (Santa Cruz Biotechnology; #sc-2357) as the secondary antibody at 1:10,000 dilution. Following each antibody incubation time, the membrane was washed 5 times with TBST for 5 min per wash. The Amersham ECL Prime substrate mixture (1:1) was applied to the membrane and incubated for 5 min before imaging with the iBright Imaging System (Invitrogen) according to the manufacturer’s protocol.

### Lysine acetylation assays

Lysine acetylation assays were performed, as previously described (23). Reactions (50 µL) contained 6 µM HvPat2 enzyme, 2 µM recombinant GK substrate, 0.2 mM acetyl-CoA, 50 mM HEPES (pH 7.5), 2 M NaCl, and 1 mM DTT. Reactions were incubated at 37 °C for 3 h and subsequently precipitated overnight on ice by the addition of 10 % (w/v) trichloroacetic acid (TCA). Samples were centrifuged (10 min, 17,999 × g, 4 °C), and the resulting protein pellets were washed twice with 100 % cold acetone. Air-dried pellets were resuspended in 20 µL 2× SDS-PAGE reducing buffer, boiled for 5 min, and analyzed by 12 % SDS-PAGE followed by CB staining and immunoblotting with anti-acetyllysine antibodies, as described above.

### Carbon shift assay

*H. volcanii* KM03 (Δ*glpK* Δ*larC*) carrying plasmids encoding His-GK ‘wild type’, K153Q, and K153R enzymes and the empty vector (pJAM202c) were used for carbon shift assays. Strains were first streaked onto glucose minimal medium (GluMM) plates and incubated at 42 °C. Single colonies were used to inoculate 5 mL GluMM liquid cultures, which were grown to early exponential phase. These cultures were then used to inoculate 50 mL of fresh GluMM at an initial OD₆₀₀ of 0.02, with incubation at 42 °C under aerobic conditions. Cells were incubated in GluMM for four days, with OD₆₀₀ measurements and sample collection performed every 24 h. At each time point, 0.5 OD₆₀₀ units of cells were harvested by centrifugation and stored at -80 °C for further analysis. On the fourth day, the remaining culture was harvested, washed three times with 18 % saltwater solution, and resuspended into GlyMM at an OD₆₀₀ of 0.02. Cells were then incubated in GlyMM for an additional four days, with OD₆₀₀ measurements and sample collection as described above. Collected samples were analyzed by SDS-PAGE followed by CB staining. Immunoblotting was performed with anti-His-tag antibodies to assess His-GK protein abundance and anti-acetyllysine antibodies to evaluate lysine acetylation profiles. As a control, mock shift experiments were performed by growing cells in GluMM and then shifting to the same carbon source (GluMM).

### Post-assay viability plating

Following growth monitoring, 5 µL from each flask culture or 96-well culture were spotted onto ATCC974, FruMM, and GlyMM agar plates. Samples were also serially diluted (1:10) in GlyMM prior to spotting. Plates were incubated at 42 °C for 5 days and imaged using the iBright FL1000 Imaging System (Thermo Fisher Scientific) in universal imaging mode to assess cell viability.

### Protein preparation and trypsin digestion for AQUA MS analysis

GK enzymes for AQUA MS analysis were purified from *H. volcanii* KM01 carrying pJAM4351 using Ni-NTA resin, as described in the previous section. Prior to purification, the cells were grown to stationary phase (OD>1) in different carbon sources including: GlyMM, GluMM, FruMM, ATCC, Gly+GluMM and Gly+FruMM. Protein concentration was determined by Bradford assay with bovine serum albumin as the standard. Purified His-GK (50 µg) was denatured in 8 M urea and 50 mM Tris-HCl (pH 8.0) containing 5 mM DTT and incubated at 37 °C for 1 h. Alkylation was performed by adding iodoacetamide to a final concentration of 15 mM, followed by incubation in the dark at room temperature for 30 min. Samples were diluted fourfold with 50 mM Tris-HCl (pH 8.0) to reduce the urea concentration to 2 M and digested overnight at 37 °C with trypsin (Promega, Madison, WI, USA) at an enzyme-to-protein ratio of 1:50 (w/w). Tryptic peptides were desalted using C18 ZipTip pipette tips (MilliporeSigma, Burlington, MA, USA) according to the manufacturer’s instructions. Briefly, ZipTips were pre-wetted with 50 % acetonitrile, equilibrated with 0.1 % formic acid, and peptides were bound to the resin. After washing with 0.1 % formic acid, peptides were eluted with 80 % acetonitrile in 0.1 % formic acid. Eluted peptides were dried using a Vacufuge Plus vacuum concentrator (Eppendorf, Hamburg, Germany) operated in V-AL mode for 45 min at 30 °C and immediately analyzed.

### Absolute quantification of lysine acetylation by AQUA-MS

Absolute quantification (AQUA) of K153 acetylation in GK was performed under different carbon source conditions using a targeted mass spectrometry approach. The Ni-NTA-purified GK samples were subjected to tryptic digestion, and synthetic heavy-isotope–labeled peptides representing the acetylated (AEWLLDNSDPIK(ac)LQR) and non-acetylated (AEWLLDNSDPIK) forms of the K153-containing tryptic peptide were used as external standards (Thermo Scientific, Rockford, IL, USA). Parallel Reaction Monitoring (PRM) analysis was performed on an Orbitrap Exploris 240 mass spectrometer coupled to a Thermo Vanquish Neo UHPLC system. Peptide separation was achieved using a 60-min LC gradient at a flow rate of 300 nL/min: 2–4 % solvent B (0.1 % formic acid in 80 % acetonitrile) in 0.5 min, 4–28 % B over 40 min, 28–42 % B over 8 min, 42– 55 % B in 4 min, followed by a final wash with 99 % B for 7 min. The Orbitrap was operated at resolving powers of 120,000 (MS1) and 30,000 (MS2) at *m/z* 200 to monitor both the +2 charged light (m/z of 700.8564 and 920.4836) and heavy peptides (m/z of 704.8635 and 925.4877) mentioned above. Quantitative data analysis was performed using Skyline (MacCoss Lab Software) (24). Chromatographic peak areas of both endogenous and heavy peptides were extracted, and quantification was based on a standard curve generated from heavy peptide dilutions (200, 1000, 2000, 10,000, and 20,000 pg). Background signal was subtracted, and all peptide abundances were normalized to their respective spiked heavy peptides (2000 pg each). Linear regression (*R²* > 0.99) confirmed high-quality quantification. The MS2 spectrum used for quantification corresponded to the y₈ ion at m/z 920.48 for the acetylated peptide (precursor m/z 998.56) and m/z 700.85 for the non-acetylated form (precursor m/z 901.46). Acetylation occupancy was calculated as the ratio of the endogenous acetylated peptide to the total of acetylated and non-acetylated peptides.

### Glycerol kinase activity assay

GK (ATP: glycerol 3-phosphotransferase, EC 2.7.1.30) enzymatic activity was measured in a coupled photometric assay, as previously described (25). GK activity was coupled to the formation of α-glycerophosphate, which was quantified in the presence of NAD^+^ and commercially available α-glycerophosphate dehydrogenase. The primary reaction mixture (1 mL) contained purified GK (0.84 μg), 3.5 mM MgCl₂, 3.5 mM ATP, 4.6 mM glycerol, 50 mM HEPES (pH 8), and 0.1 M NaCl. Reactions were incubated for 1 h at 57 °C and terminated by adding an equal volume of 0.2 N phosphoric acid (H₃PO₄), followed by centrifugation (10 min, 12,000 × g). Glycerol-3-phosphate (G3P) levels were determined enzymatically as previously described (26). An aliquot of the terminated reaction (110 µl) was mixed in a total reaction volume of 1 ml containing 0.011 N NaOH for neutralization, 1.1 mM NAD+, 0.66 M hydrazine sulfate adjusted to pH 9.4 with NaOH, 1% (w/v) nicotinamide-sodium carbonate buffer, and 8 U of rabbit muscle α-glycerophosphate dehydrogenase (EC 1.1.1.8; Sigma, 500 U). After 1 h incubation at 30°C, NADH production was measured at 340 nm. G3P concentrations were calculated by linear regression analysis (R² > 0.99) using standards linear between 0 and 5 mM G3P.

### Size-exclusion chromatography

Purified GK samples were dialyzed into 50 mM HEPES (pH 7.5), 2 M NaCl, 1 mM DTT, and 10 % glycerol. Proteins (100 µL) were loaded onto Superdex 200 HR 10/300 GL columns at 0.3 mL/min. Molecular masses were estimated using standard curves generated from known molecular weight standards: Thyroglobulin (bovine) (670 kDa), γ-globulin (bovine) (158 kDa), ovalbumin (chicken) (44 kDa), myoglobin (horse) (17 kDa), and vitamin B12 (1.35 kDa) (Bio-Rad Laboratories).

### Differential scanning fluorimetry (DSF) thermal shift assay

Thermal shift assays were performed to assess the impact of ligand binding on the stability of His-GK K153Q and K153R variants compared to wild type (wt). The proteins were purified from *H. volcanii* KM03 strains grown in GlyMM. DSF experiments were conducted as previously described (25). Briefly, each reaction contained 3 µM monomeric enzyme in a final volume of 25 µL, prepared in buffer supplemented with various combinations of 35 mM MgCl₂, 46 mM glycerol, and 35 mM ATP. SYPRO Orange protein gel stain (Invitrogen, Cat. No. S6650) was added to a final concentration of 2.5×. Enzyme-minus controls were used to correct for background fluorescence. Thermal unfolding was monitored using a C1000 Touch Thermal Cycler coupled to a CFX96 Real-Time PCR Detection System (Bio-Rad Laboratories). Samples were subjected to a linear temperature gradient from 20 °C to 95 °C at a rate of 1 °C per min. Fluorescence was recorded continuously and melting temperatures (T_m_) were determined by identifying the peak of the first derivative of the fluorescence intensity with respect to temperature. Comparisons across different ligand conditions were made to evaluate their effects on the thermal stability of the His-GK K153Q and K153R variant proteins.

### Kinetic Analysis of GK Variants

Kinetic assays were conducted using purified His-GK variants: wild-type (wt), K153Q, and K153R, expressed in the *H. volcanii* KM03 strain (Δ*glpK* Δ*larC*). Reactions were performed under optimized conditions: 100 mM NaCl, pH 8.0 (HEPES buffer), and 57 °C. GK activity was measured using the coupled enzymatic assay that monitored NADH oxidation at 340 nm. Substrate concentrations were varied to assess their effect on enzyme kinetics: glycerol (0, 0.2, 0.4, 0.6, 0.8, 1, 2, 3, and 4.6 mM), and ATP (0, 0.2, 0.4, 0.6, 0.8, 1, 2, 3, 3.5, 4, 4.5, 5, 5.5, 6, 6.5 and 7 mM). Kinetic parameters (*K_m_, K_d_, V_max_, Kc_at_, K_cat_/ K_m_*, and Hill coefficient *n*) were calculated by nonlinear regression fitting to the Michaelis-Menten, Lineweaver-Burk, and Hill equations using Excel Solver for iterative optimization. For the K153R variant with ATP, both sigmoidal and Michaelis-Menten models were tested to determine the best fit. All assays were performed in three experimental replicates, and results are presented as mean ± standard deviation.

### Growth plate assay to assess the impact of GK lysine acetylation on carbon source utilization

To evaluate the impact of GK lysine acetylation on carbon source utilization, KM06 (H1207 Δ*glpK* Δ*larC Δpat2*) was used as a host. KM06 strains were generated through transformation to individually carry the following plasmids: pJAM202c (empty vector), pJAM4351 (His-GK), pJAM4354 (His-GK K153Q acetylation mimic), and pJAM4355 (His-GK K153R non-acetylated variant). The strains were cultivated in ATCC 974 supplemented with novobiocin to stationary phase and stored in 20 % (w/v) glycerol stocks at -80 °C. For assay, strains were inoculated from the freezer stocks using a loop onto GlyMM or GluMM plates. Plates were incubated at 42 °C for 8 days. Growth was monitored by streak/colony formation. Growth on GlyMM was used as an indicator of functional complementation by the respective His-GK variants in the absence of the *pat2* gene.

### Growth curve analysis

*H. volcanii* KM06 strains, carrying pJAM202c (empty vector), pJAM4351 (His-GK), pJAM4354 (His-GK K153Q acetylation mimic), and pJAM4355 (His-GK K153R non-acetylated variant), were freshly streaked from glycerol stocks onto GluMM 0.3 µg/mL novobiocin for plasmid selection and incubated at 42°C for 5 days. A single colony from each strain was used to inoculate 5 mL of GluMM with 0.3 µg/mL novobiocin, followed by incubation at 42°C with gentle shaking (∼60 rpm) for 48 h. After incubation, culture density was measured at OD₆₀₀ using a GENESYS 40/50 Vis/UV-Vis spectrophotometer, with 1.0 OD₆₀₀ equivalent to approximately 10⁹ CFU/mL. Due to predicted sensitivity to glycerol, cells were not pre-cultured in glycerol minimal medium (GlyMM). Instead, cultures were centrifuged at 6,000 × g for 8 min, washed twice with 18% (w/v) salt water (SW), and resuspended in GlyMM. Cultures were adjusted to OD₆₀₀ 0.02, gently mixed for 15 min at room temperature, and 200 µL aliquots were transferred to each well of a 96-well plate (n = 9 per condition). To minimize edge effects, outer wells contained 300 µL of sterile nanopure water. Plates were sealed with ½-inch 3M Micropore tape. Growth was monitored at OD₆₀₀ every 15 min over 120 h at 42 °C using a BioTek Epoch 2 microplate reader (Agilent Technologies) with continuous orbital shaking. OD₆₀₀ values were normalized by subtracting the average blank signal.

### Analysis of GK abundance and acetylation in acetyltransferase and deacetylase mutant strains

To assess the role of different acetyltransferases or deacetylases in the abundance and acetylation state of GK, *H. volcanii* strains lacking acetyltransferase or deacetylase genes were generated and analyzed. Strains KM05 (Δ*pat1*), KM06 (Δ*pat2*), KM07 (Δ*sir2*), and KM08 (Δ*elp3*) were each transformed with plasmids expressing His-GK or the empty vector pJAM202c as a control. Cultures were grown in ATCC, GluMM, FruMM, and GlyMM media at 42 °C with orbital shaking (200 rpm) to stationary phase (OD₆₀₀ > 1.0). Aliquots corresponding to 0.5 OD₆₀₀ units were harvested by centrifugation, and cell pellets were stored at −80 °C until further analysis. For protein analysis, pellets were resuspended directly in 2× Laemmli sample buffer, boiled for 5 min, and centrifuged at RT (10 min, 13,000 × g). The equivalent of 0.1 OD of each sample was loaded onto the SDS-PAGE gels. Proteins were resolved by SDS-PAGE and analyzed by CB staining, as well as immnoblotting using anti-His and anti-acetyllysine (Kac) antibodies to assess His-GK abundance and acetylation status. This experimental setup enabled direct evaluation of the impact of acetyltransferase and deacetylase deletions on His-GK expression and acetylation *in vivo*.

### Statistical Analysis

All experiments included technical replicates performed in at least triplicate and were independently repeated at least three times to ensure reproducibility. Data are presented as mean values ± standard deviation (SD). Statistical significance between experimental groups was determined using Student’s *t*-test, with *p* < 0.05 considered statistically significant. All statistical analyses were performed using Microsoft Excel.

## Data availability

The MS-based proteomic datasets generated in this study were deposited in the UCSD MassIVE repository (Mass Spectrometry Interactive Virtual Environment, https://massive.ucsd.edu) under the following accession numbers ID: MSV000097918, MSV000097919, MSV000097920 and MSV000097921 (protected until publication with the password: maupinlab).

## RESULTS

### Lysine acetylation of *H. volcanii* GK

To provide deeper coverage of the lysine acetylation sites of *H. volcanii* GK, the purified enzyme was analyzed by LC-MS/MS. Compared to the single K153 acetylation site identified through MS-based proteomic analysis of cells grown in glycerol medium (9), two sites of acetylation (K121 and K153) were observed when the GK enzyme was analyzed after purification from glycerol-grown cells. To better understand the conservation of these sites and how lysine acetylation may impact the structure and function of GK, 3D-modeling was used to map the location of K153 and K121 in relationship to the binding sites for the substrates ATP and glycerol **(Fig. 1A).** Haloarchaeal proteins possess highly acidic surfaces compared to their mesohalic counterparts, an adaptation that supports protein function within the high-salt cytosol of organisms that employ a ‘salt-in’ strategy (27). Surface electrostatic modeling predicted K121 to be in a highly acidic region on the GK surface, with the basic charge of K121 appearing to have limited impact on the overall charge in this region (**Fig. 1B**). By contrast, K153 was found to reside in a predominantly positive/neutral region of the 3D-model, suggesting potential accessibility for protein interactions **(Fig. 1C).** Examination of the GK structure further indicated that K153 is embedded within a flexible loop motif **(Fig. 1D).** Sequence alignment of GK homologs across the *Halobacteria* class revealed that K153 is highly conserved, with K121 also conserved but to a lesser extent (**Fig. 1E**). Further examination revealed K153 and the flanking flexible loop are absent in bacterial and eukaryotic counterparts. These observations suggest a unique adaptation and potentially functional role for K153 acetylation in *Halobacteria* GKs.

**Fig 1.**
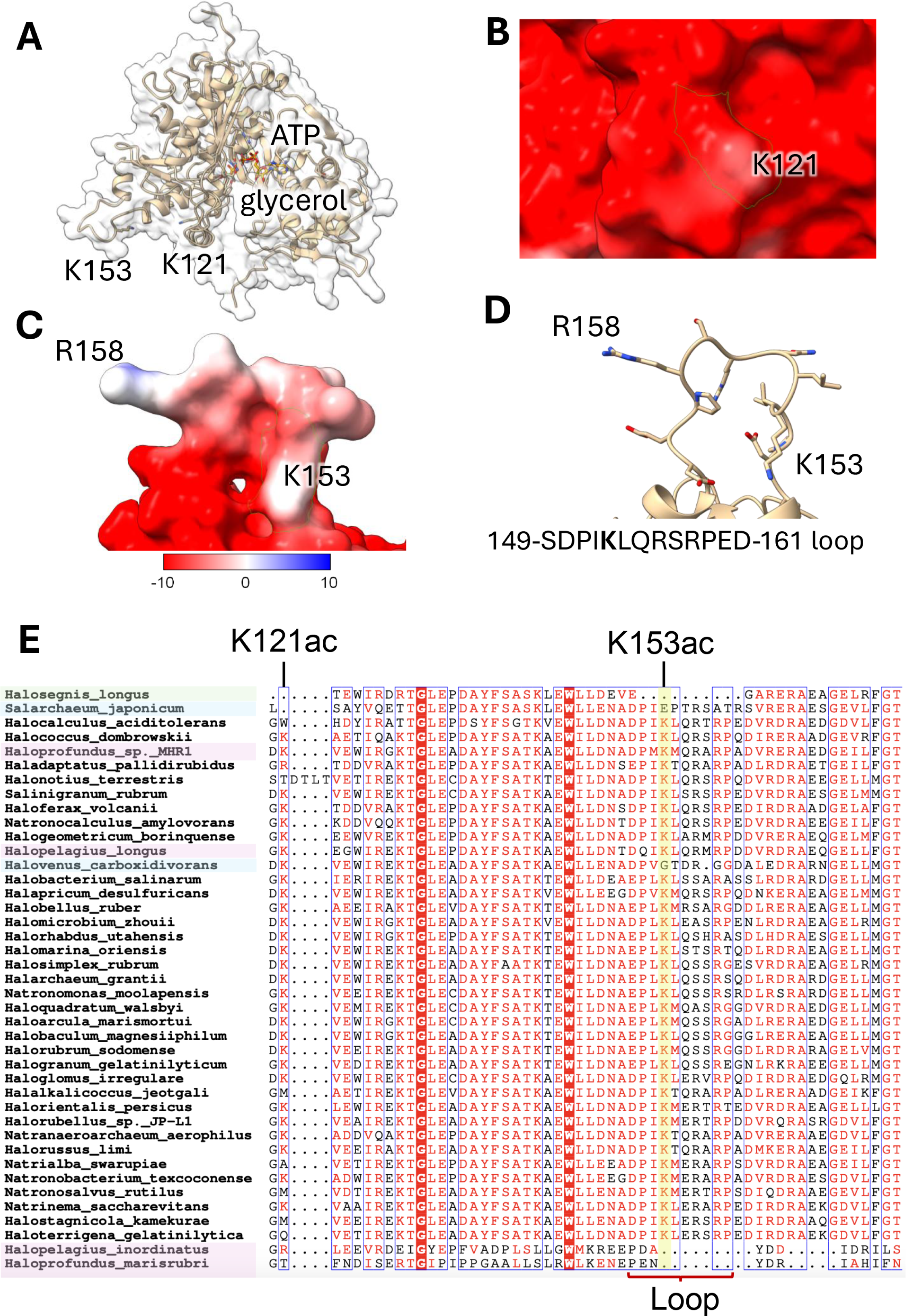
Structural and evolutionary features of lysine acetylation sites in *H. volcanii* GK. A. Predicted 3D-structure of GK with mapped functional sites. AlphaFold2-predicted structure of *H. volcanii* GK showing ATP and glycerol binding sites. Lysine residues K121 and K153, identified as acetylated by LC-MS/MS in glycerol-grown cells, are indicated. B. Electrostatic surface representation showing K121 (outlined in green) located within a predicted acidic region on the surface of GK. C. Electrostatic surface representation showing K153 (outlined in green) positioned in a predicted surface-exposed loop within a region of neutral to positive charge. D. Structural context of K153 within a predicted flexible loop. K153 is located within a surface-exposed loop predicted to be flexible based on AlphaFold2 confidence metrics and structural features. E. Conservation of the K153 region among *Halobacteria*. Multiple sequence alignment of GK homologs from the *Halobacteria* class (**Dataset S1**) shows that K153 and the surrounding loop region are highly conserved among haloarchaea.

### K153 is the major lysine acetylation site of *H. volcanii* GK

To assess the contribution of K153 acetylation on GK regulation, GK enzymes with substitutions that mimicked the acetylated (K153Q) and non-acetylated (K153R) states were generated. The wild type (wt) and variant GK proteins were expressed with N-terminal His tags from plasmids and purified by Ni-NTA enrichment from *H. volcanii* H1207 strains grown on ATCC 974 rich medium. SDS-PAGE and anti-His immunoblotting revealed comparable expression and purification yields of the GK proteins across all constructs **(Fig. 2AB).** Immunoblotting using anti-acetyllysine antibodies demonstrated a strong acetylation signal in the eluate of His-GK wt compared to the K153Q and K153R variants **(Fig. 2C)**. These data indicate that K153 is the predominant acetylation site of the *H. volcanii* GK under standard growth conditions (in rich medium with yeast extract and tryptone). To independently validate the specificity of K153 acetylation, *in vitro* acetylation assays were performed using the acetyltransferase Pat2, previously reported to acetylate *H. volcanii* GK (23). MS data confirmed the identity and deacetylated state of the recombinant *H. volcanii* His-GK produced in *E. coli* Rosetta DE3. The non-acetylated GK wt and variant (K53Q and K153R) proteins prepared from recombinant *E. coli* **(Fig. 3A)** were incubated with Pat2 using acetyl-CoA as the acetyl-group donor. Lysine acetylation was assessed by immunoblotting. Consistent with *in vivo* observations, only the GK wt enzyme was acetylated by Pat2, whereas the K153Q and K153R variants remained unmodified **(Fig. 3B).** These results show that K153 is the primary target of Pat2-mediated acetylation of *H. volcanii* GK.

**Fig 2.**
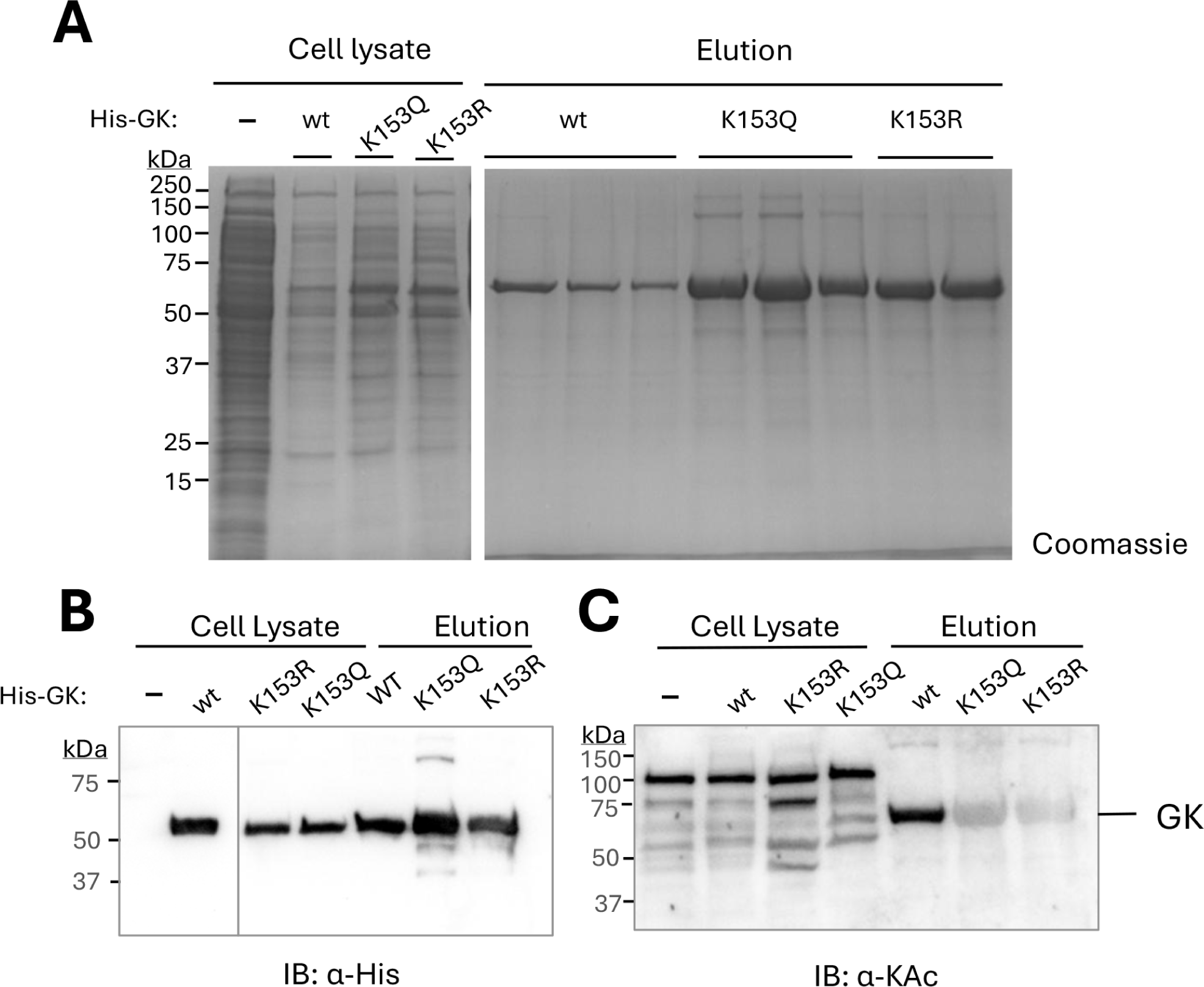
Detection of K153 as the primary acetylation site in *H. volcanii* GK. A. Purification of GK enzymes. *H. volcanii* H1207 strains carrying plasmids expressing His-GK wt, K153Q and K153R or an empty vector control (–) were grown to stationary phase in ATCC 974 medium. Cells were lysed (cell lysate) and proteins were purified by Ni-NTA resin (elution). Protein samples were separated by 12 % SDS-PAGE and stained with CB. B. Detection of His-tagged GK proteins. Samples analyzed by immunoblotting using anti-His antibodies (IB: α-His) to assess the abundance of His-tagged GK enzymes in the elution fractions. C. Analysis of lysine acetylated proteins. Protein samples were analyzed by immunoblotting using anti-acetyllysine antibodies (IB: α-KAc) to compare the lysine acetylated state of proteins in the cell lysate and Ni-NTA elution fractions. All experiments were performed in biological triplicates. Representative immunoblots are shown.

**Fig 3.**
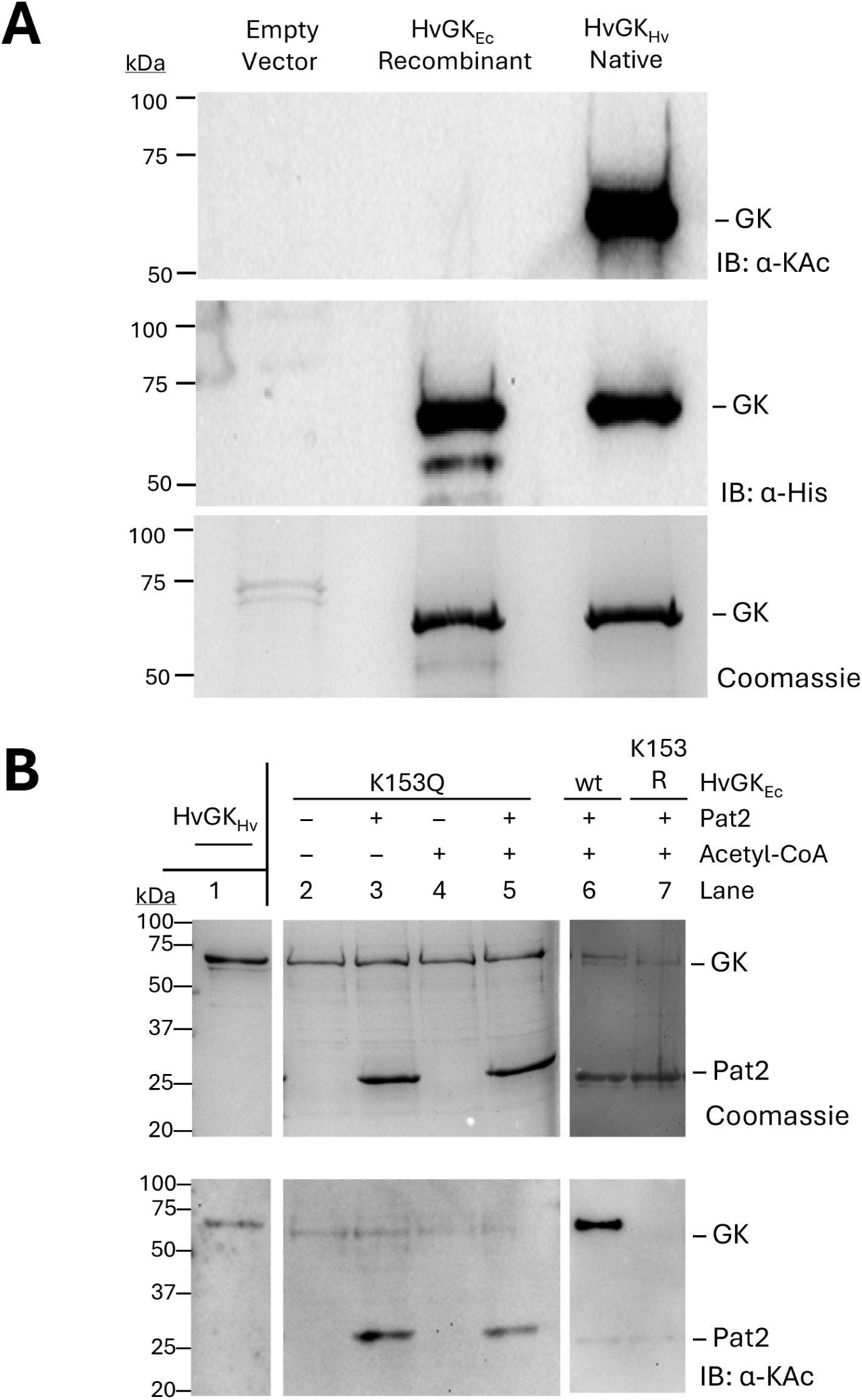
Pat2 selectively acetylates the K153 site of *H. volcanii* GK. A. Absence of the lysine acetylation of *H. volcanii* GK when expressed and purified from recombinant *E. coli*. Proteins were purified by Ni-NTA chromatography from *H. volcanii* H1207 strains expressing His-GK from a plasmid (HvGK_Hv_) or the empty vector control compared to recombinant *E. coli* expressing the His-GK enzyme (HvGK_Ec_). Total protein abundance in sample was detected by CB staining. His-GK enzyme was detected by IB: α-His. Lysine acetylation levels were assessed by IB: α-KAc. B. *In vitro* acetylation of HvGK_Ec_ wt compared to the K153 variant proteins. Reactions included 2 µM of HvGK_Ec_ (wt, K153Q or K153R) incubated with 6 µM Pat2 acetyltransferase and 0.2 mM acetyl-CoA, as indicated. Acetylation was detected by IB: α-KAc. CB staining confirmed equal protein loading. HvGK_Hv_ was directly applied to the SDS-PAGE gels to compare its migration to the HvGK_Ec_ enzymes. All experiments were performed in biological triplicates. Representative immunoblots are shown.

### Carbon source modulates GK acetylation *in vivo*

Given the importance of carbon source availability in regulating central metabolism, we investigated whether carbon source conditions influenced GK acetylation. Initially, a contaminant band was observed in the elution fraction of strain H1207 harboring the empty vector control when grown in glucose minimal medium (GluMM), migrating close to the expected molecular weight of GK. Mass spectrometry analysis identified this band as LarC, a putative nickel insertion protein harboring a native histidine-rich domain **(Fig. S1).** To eliminate interference from LarC, a *glpK* and *larC* double mutant (KM03) was constructed to serve as the host strain for the carbon source experiments. The N-terminal His-GK wt and variant (K153Q, K153R) enzymes were expressed from plasmids in *H. volcanii* KM03 strains cultured in minimal media with glycerol or glucose as the sole carbon source (GlyMM or GluMM). Proteins were purified, and cell lysates (input) and elution fractions (output) were analyzed by SDS-PAGE followed by anti-His and acetyl-lysine immunoblotting **(Fig. 4).** The wt His-GK enzyme displayed low lysine acetylation levels in glucose-grown cells (*lane 6*), with a pronounced increase in acetylation observed when cells were grown on glycerol as the carbon source (*lane 14*) (**Fig. 4**). In contrast, the His-GK variants carrying K153Q (*lanes 7 and 15*) or K153R (*lanes 8 and 16*) substitutions showed no detectable acetylation under these conditions (**Fig. 4**), consistent with observations in cells grown in ATCC 974 medium (**Fig. 2C**). Overall, these results suggest that GK acetylation is upregulated in cells grown on glycerol, and reveal carbon source-dependent modulation of acetylation status.

**Fig 4.**
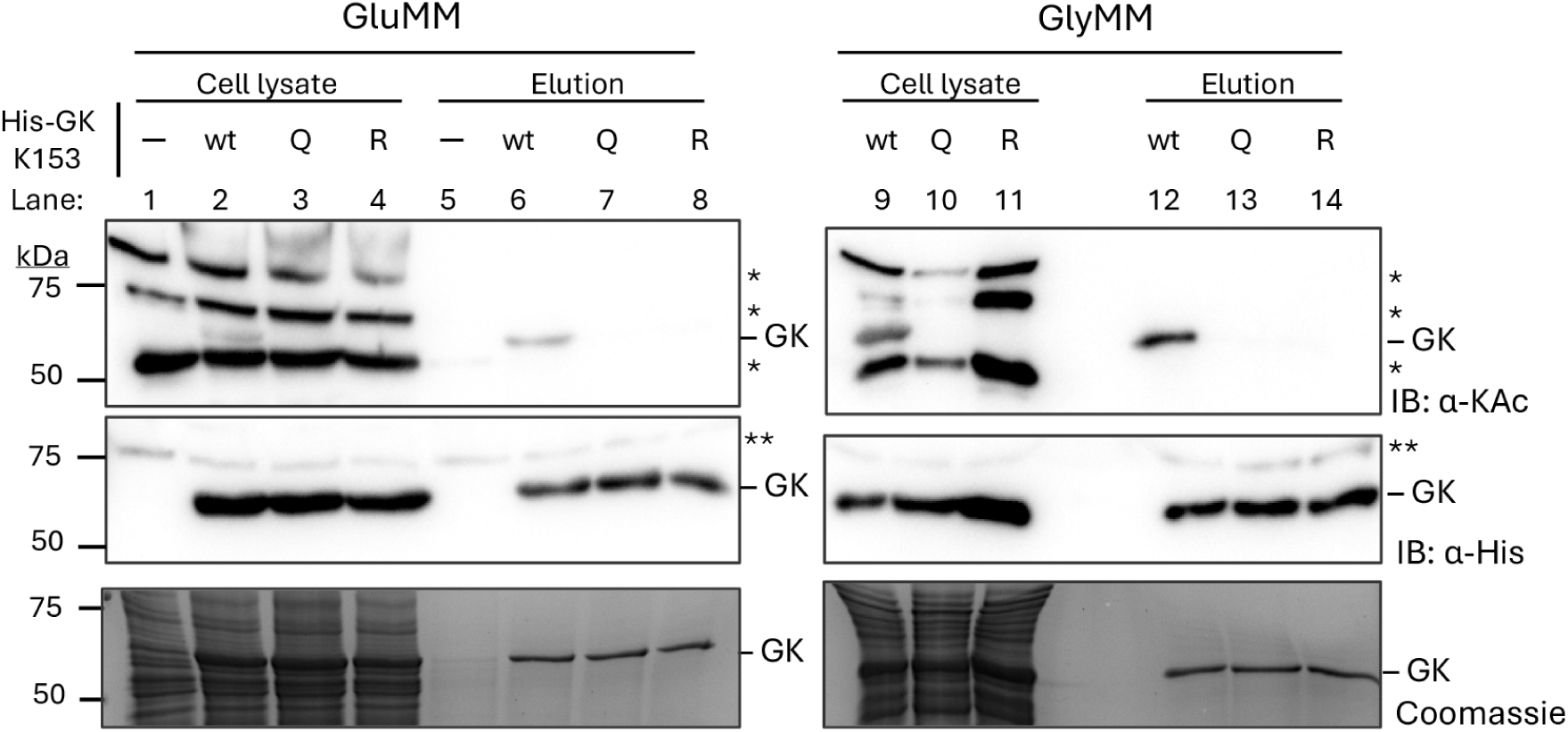
K153-acetylated GK, the major acetylated form of this enzyme, is elevated in glycerol-vs. glucose-grown cells. His-GK and variant proteins were purified by Ni-NTA affinity chromatography from *H. volcanii* KM03 (Δ*glpK* Δ*larC*) strains expressing His-GK wt, K153Q or K153R that were grown in GluMM or GlyMM, as indicated. GluMM-grown KM03 carrying the empty vector (pJAM202c, indicated by ─) was included as a control; this strain did not grow in GlyMM due to the essential role of GK under these conditions. IB: α-His and CB staining were used to determine the abundance of the His-tagged proteins and confirm equal protein loading, respectively. IB: α-KAc was used to examined lysine acetylation. Clarified lysates (Ni-NTA input) were diluted 1:10, and all elution fractions were normalized to 0.5 µg protein. Theoretical molecular weight of GK is 56.7 kDa. Asterisks: (*) lysine acetylated proteins detected in cell lysate in addition to GK; (**) non-specific signal detected with anti-His antibody in empty vector control. All experiments were performed in biological triplicates. Representative immunoblots are shown.

### Carbon shift reveals dynamic acetylation regulation and functional consequences

To further explore the dynamics of GK acetylation during carbon source transitions, carbon shift experiments were conducted. *H. volcanii* KM03 strains expressing His-GK wt, K153Q, and K153R from plasmids, or carrying the empty vector, were grown for four days in GluMM, followed by a shift to GlyMM for an additional four days. Cell samples were collected daily to monitor acetylation status and growth (OD₆₀₀ measurements). Immunoblot analysis showed that the lysine acetylation levels of GK increased following the shift to glycerol medium **(Fig. 5A).** In contrast, no acetylation of GK was detected in the strains expressing the His-GK K153Q or K153R variants or the empty vector control throughout the experiment. Growth curve analysis revealed that cells expressing His-GK wt or K153Q exhibited continued growth after the shift in carbon source. At the same time, the K153R-expressing and empty vector strains experienced growth arrest **(Fig. 5B and C**). Control assays, in which cells were transferred from GluMM to GluMM, showed no significant growth differences among the strains (**Fig. 5C**), indicating that the growth defect observed in KM03 strains lacking GK (empty vector) or expressing the K153R deacetylation mimic was specific to the carbon source shift from glucose to glycerol. SDS-PAGE and anti-His immunoblotting confirmed the His-GK wt, K153Q and K153R proteins were expressed under these control assay conditions **(Fig. S2).** Notably, anti-acetyllysine immunoblots showed a gradual decrease in overall lysine acetylation levels across all strains during extended growth in GluMM (**Fig. S2** lanes 6 to 8) This trend suggests that lysine acetylation in *H. volcanii* is responsive to carbon source conditions and may be broadly shifted to alternative substrate proteins in glucose-rich environments. These results support the idea that dynamic acetylation, beyond GK alone, plays a role in metabolic adaptation and may contribute to the distinct growth responses observed upon transitioning to glycerol.

**Fig 5.**
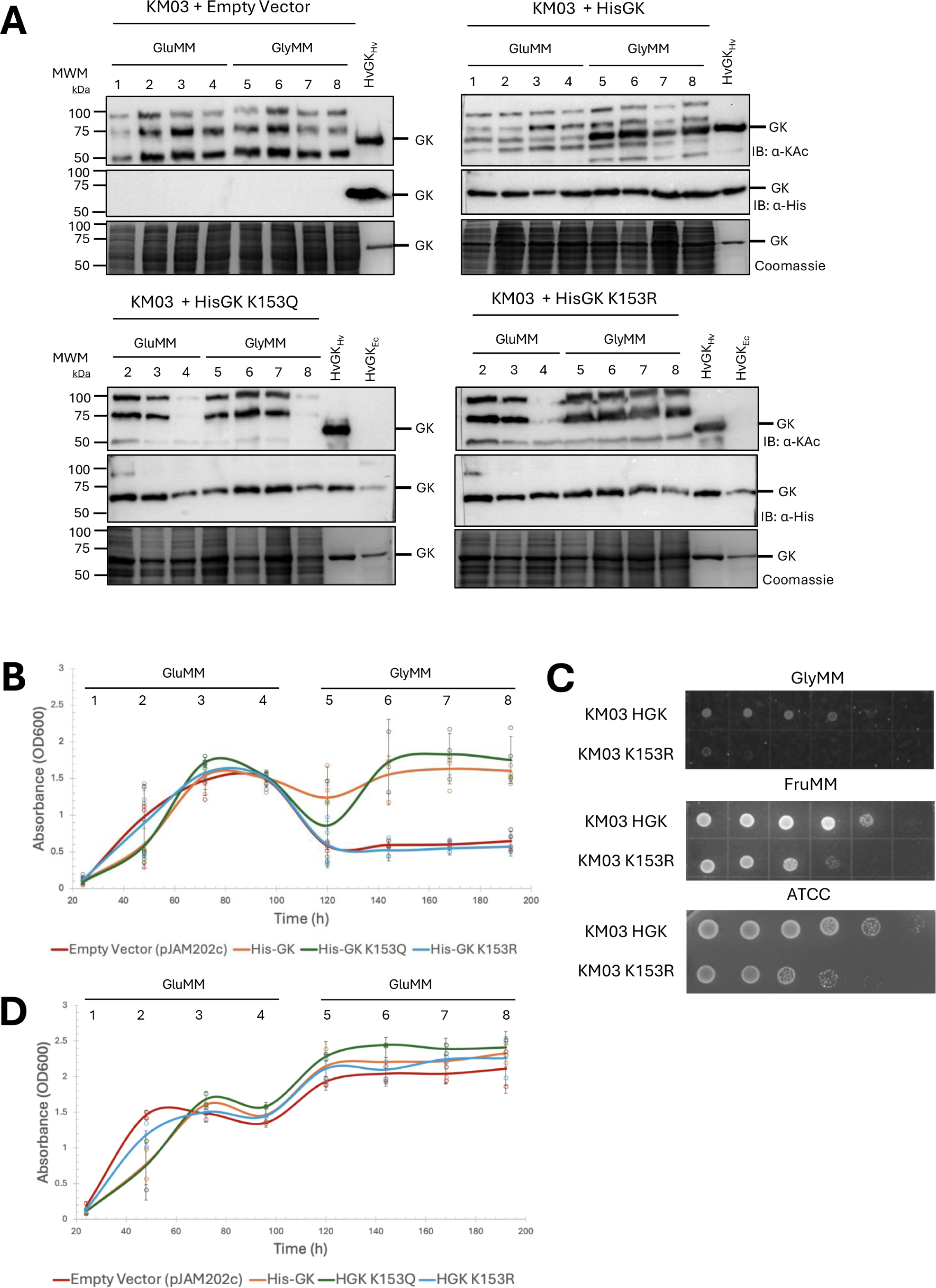
Dynamic increase in the abundance of K153-acetylated GK and its requirement for *H. volcanii* growth following shift from glucose to glycerol as the carbon source. A. Increased acetylation of His-GK after a shift from glucose to glycerol as a carbon source. Whole-cell lysate of *H. volcanii* KM03 strains expressing His-GK wt, K153Q, and K153R were compared to an empty vector control. Cells were grown in GluMM for 4 days and then shifted to GlyMM for an additional 4 days. The time course is represented with lanes numbered according to days: 1 (24 h), 2 (48 h), 3 (72 h), 4 (96 h), 5 (120 h), 6 (144 h), 7 (168 h), and 8 (192 h). Equal loading of cell lysate was confirmed by CB stain. His-GK wt and variant protein abundance and lysine acetylation status were determined by IB: α-His and α-KAc, respectively. His-GK purified from *H. volcanii* (HvGK_Hv_) and recombinant *E. coli* (HvGK_Ec_) served as positive controls for the α-His IB and as positive and negative controls, respectively, for detecting lysine-acetylated GK by α-KAc IB. B. Importance of the acetylation of GK K153 on the growth of *H. volcanii* after the shift from glucose to glycerol. KM03 strains expressing His-GK wt, K153Q, K153R, or empty vector were monitored for growth by OD₆₀₀ over eight days with the shift from GluMM to GlyMM, as indicated. C. Following growth monitoring, samples KM03 HGK wt and K153R were serially diluted and 5 µL from each strain were spotted onto GlyMM, FruMM and ATCC974 to assess cells viability. D. The mock shift from GluMM to GluMM confirms that the growth defect in the strain expressing the GK K153R variant results from the glucose-to-glycerol shift, not from culture manipulation. KM03 strains expressing His-GK wt, K153Q, K153R, or empty vector were grown in GluMM and transferred to fresh GluMM as indicated with growth monitored by OD₆₀₀ over eight days. All experiments were performed in biological triplicates. Representative immunoblots are shown.

### GK acetylation levels correlate with its catalytic activity across carbon sources

To investigate whether acetylation status correlates with GK enzymatic activity, we purified His-GK from KM01 cells grown in various carbon sources, including glycerol, fructose, glucose, and rich (ATCC 974) media. Additionally, the enzyme was analyzed when expressed in cells grown in mixed carbon sources (glycerol + glucose MM and glycerol + fructose MM). Purified His-GK proteins were evaluated by SDS-PAGE, immunoblotting using anti-His and anti-acetyl-lysine antibodies, targeted quantitative mass spectrometry (AQUA analysis), and glycerol kinase activity assays. SDS-PAGE and immunoblotting revealed distinct acetylation patterns depending on the carbon source, with the highest levels of His-GK acetylation detected in cells grown with glycerol supplementation **(Fig. 6A)**. AQUA-MS analysis demonstrated that GK K153 acetylation levels were highest in glycerol-grown samples, with the ratio of acetylated to non-acetylated peptide reaching 3.5 in GlyMM, revealing 78 % occupancy of the K153 acetylation site. Acetylation ratios decreased progressively under mixed or alternative carbon sources: 2.9 (74%) in Gly+FruMM, 2.4 (71 %) in Gly+GluMM, 1.0 (50 %) in FruMM, 0.5 (33 %) in ATCC rich medium, and 0.2 (17%) in GluMM **(Fig. 6B).** GK purified from glycerol-grown cells, which showed the highest acetylation, also displayed the highest enzymatic activity **(Fig. 6C).** Plotting activity against acetylation ratio revealed a strong positive correlation (R² = 0.90), indicating that higher acetylation levels are associated with increased GK activity **(Fig. 6D).** This suggests that acetylation enhances GK catalytic efficiency. Additionally, the acetylation mimic variant K153Q exhibited enzymatic activity comparable to wild-type His-GK, whereas the non-acetylated mimic K153R displayed significantly reduced activity relative to both wild-type and K153Q **(Fig. 6E)**. Together, these results suggest that acetylation at K153 enhances His-GK activity and is responsive to carbon source availability.

**Fig 6.**
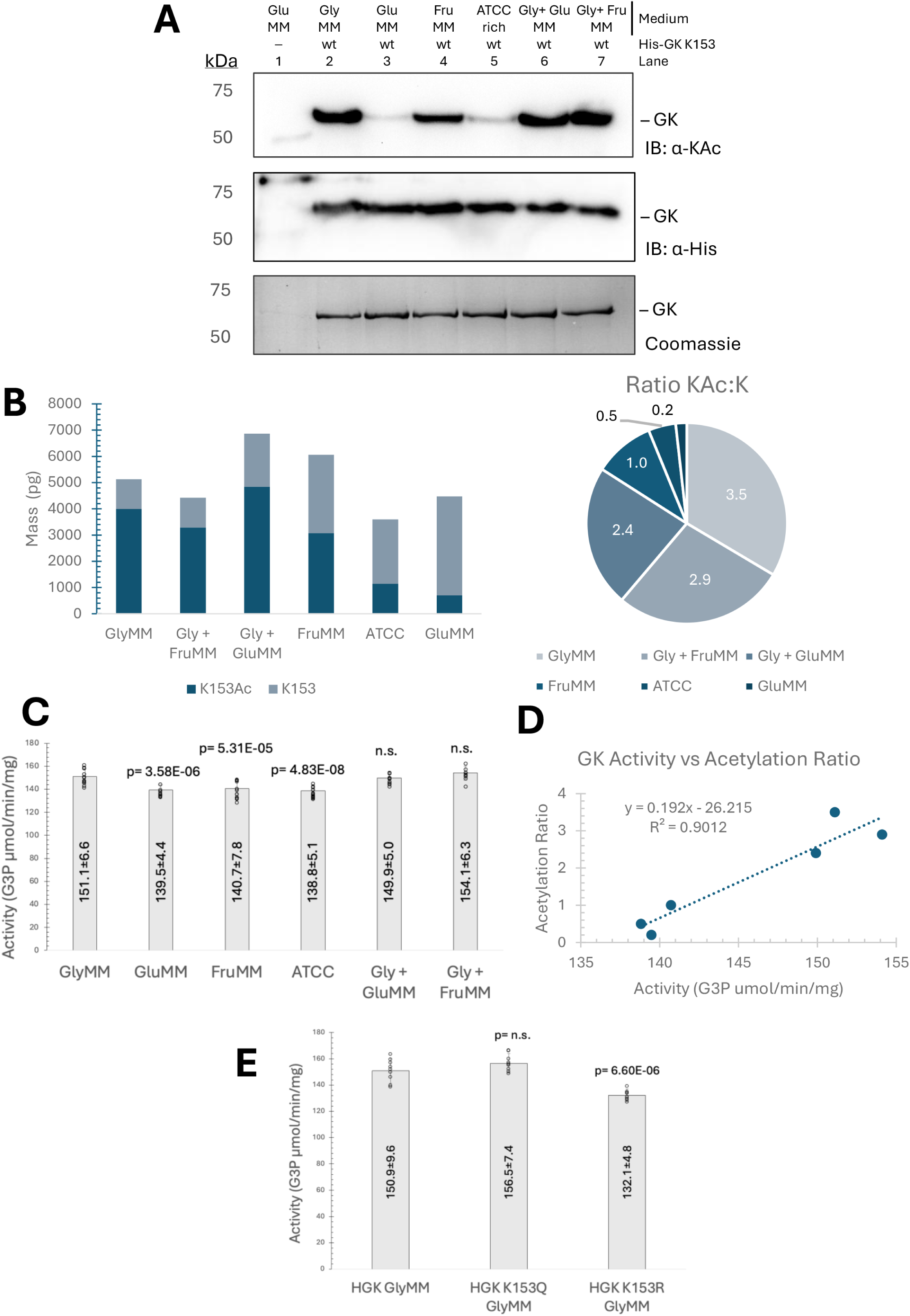
K153-acetylated GK abundance and enzymatic activity are positively correlated and vary across different carbon sources. A. Lysine acetylation status of His-GK purified from cells grown in different carbon sources. His-GK was purified by Ni-NTA resin from KM01-pJAM4351 cells grown to stationary phase in GlyMM, FruMM, GluMM, rich medium (ATCC), or mixed conditions (Gly+GluMM, Gly+FruMM). His-GK proteins were separated by SDS-PAGE and detected by CB staining and IB: α-His. Lysine acetylation levels of the His-GK proteins were analyzed by IB: α-KAc. B. Quantification of K153 acetylation occupancy by AQUA-MS. Targeted mass spectrometry was used to determine the acetylated:non-acetylated peptide ratio at K153 under each growth condition. Acetylation occupancy was highest in GlyMM (78%, 3.5:1 ratio) and decreased in mixed or non-glycerol conditions. C. GK enzyme activity varies with carbon source used for cell growth. Specific activity of His-GK purified from cells grown on the different carbon sources was measured in a coupled assay. p-values represent comparison to the His-GK enzyme purified from GlyMM. D. Correlation between acetylation levels and GK activity. GK activity (G3P µmol/min/mg) was plotted against K153 acetylation ratios from AQUA-MS analysis. A linear regression fit yielded y = 0.192x – 26.215 with R² = 0.90, indicating a strong positive correlation between acetylation and enzymatic activity. E. Activity of acetylation mimic and non-acetylated GK variants. Specific activity of His-GK wt, K153Q (acetylation mimic), and K153R (non-acetylated mimic) proteins purified from *H. volcanii* KM03 strains grown on GlyMM. p-values represent comparison to His-GK wt. Enzymatic activity values represent the mean of three independent experiments, each with three technical replicates. Statistical significance was determined using an unpaired two-tailed t-test (p < 0.05 considered significant; n.s., not significant).

### Acetylation state influences GK oligomeric conformation

Previous study reported *H. volcanii* GK predominantly exists as a homodimer but can form a homotetramer upon the presence of glycerol (3). To evaluate whether acetylation state and carbon source conditions impact *H. volcanii* GK oligomerization, we performed SEC analyses of GK purified from cells grown in GluMM, FruMM, and rich medium (ATCC). SEC profiles demonstrated that His-GK predominantly eluted as a dimer under all tested conditions, regardless of glycerol supplementation during chromatography **(Fig. 7A–C).** No tetrameric shift was observed. These findings contrast with previous observations in glycerol-grown cultures (3), suggesting that acetylation status influences substrate-induced oligomerization.

**Fig 7.**
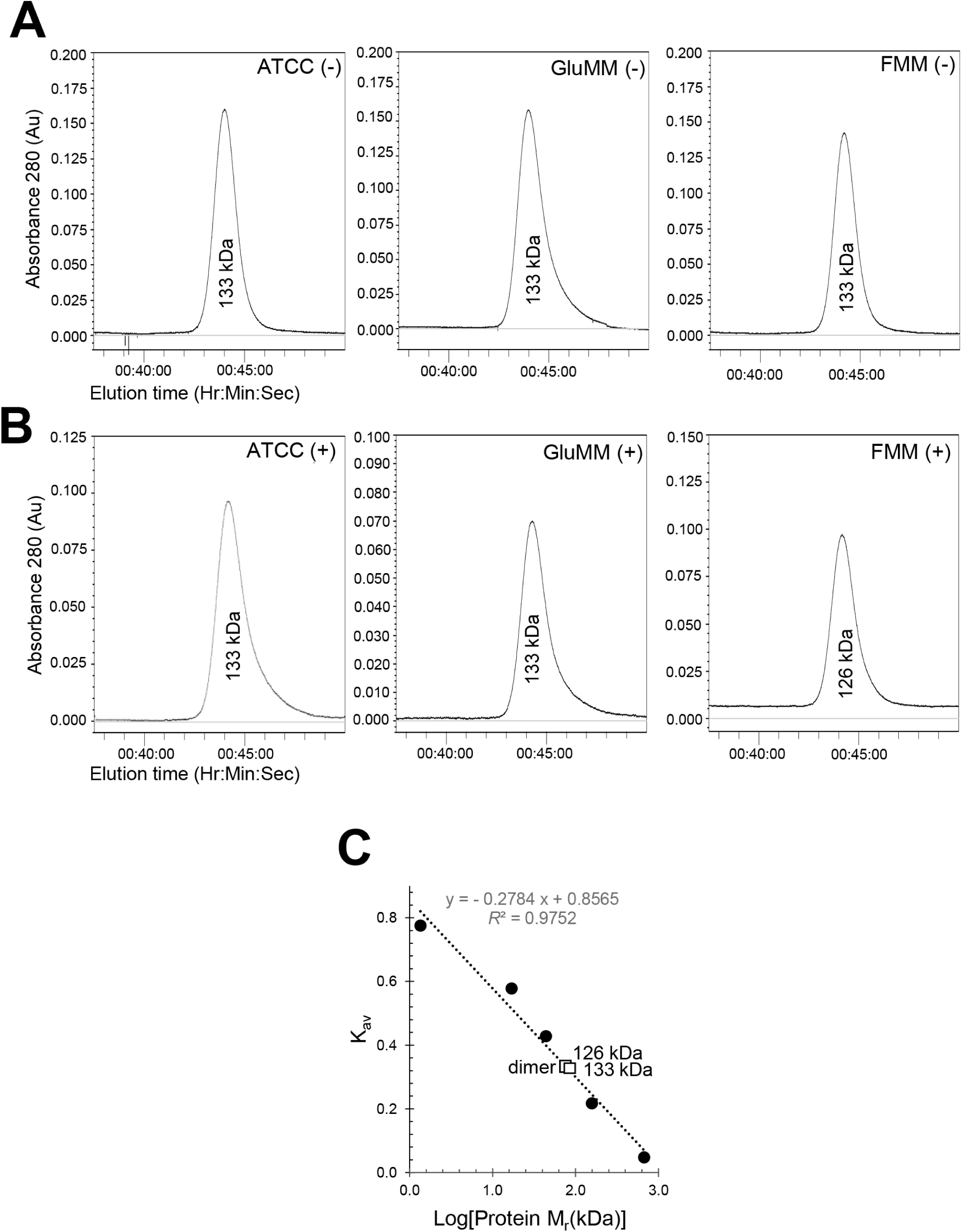
GK oligomeric state under different carbon source and buffer conditions. A. His-GK purified from *H. volcanii* cells grown in ATCC, GluMM, or FruMM elutes as a dimer. Size exclusion chromatography (SEC) profiles of His-GK purified from *H. volcanii* grown in ATCC rich medium, GluMM, or FruMM show a single predominant peak corresponding to a homodimer (∼113 kDa). No peaks corresponding to tetrameric forms were observed under these growth conditions. B. Glycerol supplementation during chromatography does not alter oligomeric state. His-GK purified from the same conditions as in panel A were analyzed by SEC in buffer supplemented with 10 % (v/v) glycerol. Elution profiles remained consistent with the dimeric state, indicating that addition of glycerol during chromatography does not promote higher-order assembly. C. SEC standard curve confirms dimeric His-GK elution profile. SEC elution positions of His-GK were compared to molecular mass standards (closed circles) using Kav (Mr) values. His-GK homodimer (open square) eluted at Kav 0.272 (126 kDa) and 0.265 (133 kDa), consistent with a dimeric state. Theoretical molecular weights: 56.7 kDa monomer and 113.4 kDa dimer. SEC was performed using Superdex 200 HR 10/300 GL columns in 50 mM HEPES (pH 7.5), 2 M NaCl, 1 mM DTT, with or without 10% glycerol.

### Impact of K153 acetylation and ligand binding on His-GK thermal stability

Thermal shift assays were performed to assess how acetylation at K153 affects the thermal stability of *H. volcanii* His-GK. The wt His-GK was compared to the two variants: K153Q, which mimics the acetylated state, and K153R, which mimics the non-acetylated form (**Table 1**). In the absence of ligands, both wt and K153Q exhibited a melting temperature (Tm) of 79 °C, whereas K153R showed significantly reduced stability, with a Tm of 77 °C compared to wt. In the presence of ATP, the K153R again had a lower thermal stability with a Tm of 78 °C when compared to the wt and K153Q enzymes which displayed a Tm of 80 °C. Similar patterns were observed in the presence of MgCl₂, ATP plus glycerol, ATP plus MgCl₂, and the combination of all three ligands, where K153R consistently showed significantly lower Tm values than wild-type. Moreover, the K153Q variant consistently mirrored or slightly exceeded wt thermal stability across the tested conditions. The exception was glycerol alone, which increased the Tm of all three enzymes to 85–86 °C, with no significant differences observed under this condition. Overall, these results indicate that acetylation at K153 enhances His-GK thermal stability and that glycerol binding provides a substantial stabilizing effect regardless of acetylation status, likely contributing to enzyme resilience under metabolically relevant conditions.

**Table 1.** Melting temperature (Tm) values as determined by differential scanning fluorimetry (DSF).

| *H. volcanii*<br>strain: | KM03-pJAM4351 | KM03-pJAM4354 | KM03-pJAM4355 |
| --- | --- | --- | --- |
| **His-GK:** | wt | K153Q | K153R |
| **KAc state:** | High Ac | Mimic Ac | Mimic no Ac |
| **Ligand(s)** | T<sub>m</sub> values (°C)<sup>a</sup> |  |  |
| ATP<br>Mg<br>Glycerol | 90 ± 0.5 | 90 ± 0.6 (n.s.) | 88 ± 0.0** |
| ATP | 80 ± 0.5 | 80 ± 0.0 (n.s.) | 78 ± 0.6** |
| Mg | 78 ± 1.0 | 78 ± 0.0 (n.s.) | 76 ± 0.6** |
| Glycerol | 85 ± 0.0 | 86 ± 0.6 (n.s.) | 86 ± 0.0 (n.s.) |
| ATP<br>Glycerol | 88 ± 0.6 | 88 ± 0.6 (n.s.) | 86 ± 0.0** |
| Mg<br>Glycerol | 84 ± 0.0 | 84 ± 0.0 (n.s.) | 85 ± 0.0 (n.s.) |
| ATP<br>Mg | 81 ± 0.6 | 81 ± 0.6 (n.s.) | 79 ± 0.6** |
| Buffer only | 79 ± 0.5 | 79 ± 0.6 (n.s.) | 77 ± 0.6** |

<sup>a</sup>T<sub>m</sub> values of His-GK wt, K153Q, and K153R enzymes were determined by DSF following purification by Ni-NTA affinity chromatography from *H. volcanii* KM03 strains grown to stationary phase in GlyMM. Assays were performed in 50 mM HEPES (pH 7.5), 2 M NaCl, and included ATP (35 mM), MgCl<sub>2</sub> (35 mM), and glycerol (46 mM) as indicated. The wt His-GK (pJAM4351) was used as the reference for statistical comparisons against K153Q (acetylation mimic; pJAM4354) and K153R (non-acetylated mimic; pJAM4355). Statistical analysis was performed using unpaired two-tailed *t*-tests. Significant differences ( $p < 0.05$ ) between wt and K153R were observed under the following conditions: no ligands ( $p = 2.63 \times 10^{-3}$ ), ATP ( $p = 4.09 \times 10^{-4}$ ), MgCl<sub>2</sub> ( $p = 0.01$ ), ATP + glycerol ( $p = 3.54 \times 10^{-3}$ ), ATP + MgCl<sub>2</sub> ( $p = 4.42 \times 10^{-3}$ ), and ATP + MgCl<sub>2</sub> + glycerol ( $p = 4.14 \times 10^{-3}$ ). No significant differences were detected between wt and K153Q under any condition. \*\* in the table indicates a significant difference relative to wt; “n.s.” = not significant. All measurements were performed in biological triplicate. Supporting data are provided in **Figs. S3–S5**.

### Acetylation at K153 modulates GK kinetics and allosteric behavior

To investigate the biochemical impact of the lower thermostability of the K153R vs. wild-type enzyme and to evaluate how acetylation at K153 affects GK activity and substrate recognition, the kinetic parameters for wild-type GK (His-GK wt), the acetylation mimic (K153Q), and the non-acetylatable mutant (K153R) were measured using glycerol and ATP as substrates. All enzymes were purified from a Δ*glpK* background (KM03) and assayed under optimal activity conditions. With glycerol as the variable substrate, both His-GK wt and K153Q displayed similar *K_d_* values (∼2 mM) and high catalytic turnover (*K_cat_* >210 s⁻¹), with Hill coefficients of 1.52 and 1.42, respectively, indicating positive cooperativity **(Fig. 8AC).** In contrast, K153R showed a nearly 2-fold reduction in *K_cat_* (∼117 s⁻¹), despite a lower apparent *K_d_* (1.08 mM), suggesting loss of catalytic efficiency. Interestingly, K153R exhibited an even higher Hill coefficient (*n* = 2.29), pointing to altered cooperative behavior possibly arising from conformational or oligomeric shifts **(Fig. 8C)**. When ATP was varied, His-GK wt and K153Q retained sigmoidal kinetics, with Hill coefficients ∼1.5 and *K_cat_* values exceeding 270 s⁻¹. In contrast, K153R displayed a markedly different profile. When fit to the Hill equation, the K153R mutant showed a modest Hill coefficient (*n* = 1.38), but the kinetic curve was more accurately described by the Michaelis– Menten model, with a *K_m_* of 4.23 mM **(Fig. 8BC)**. This shift from cooperative to non-cooperative behavior supports a role for K153 acetylation in maintaining allosteric regulation of ATP binding.

**Fig 8.**
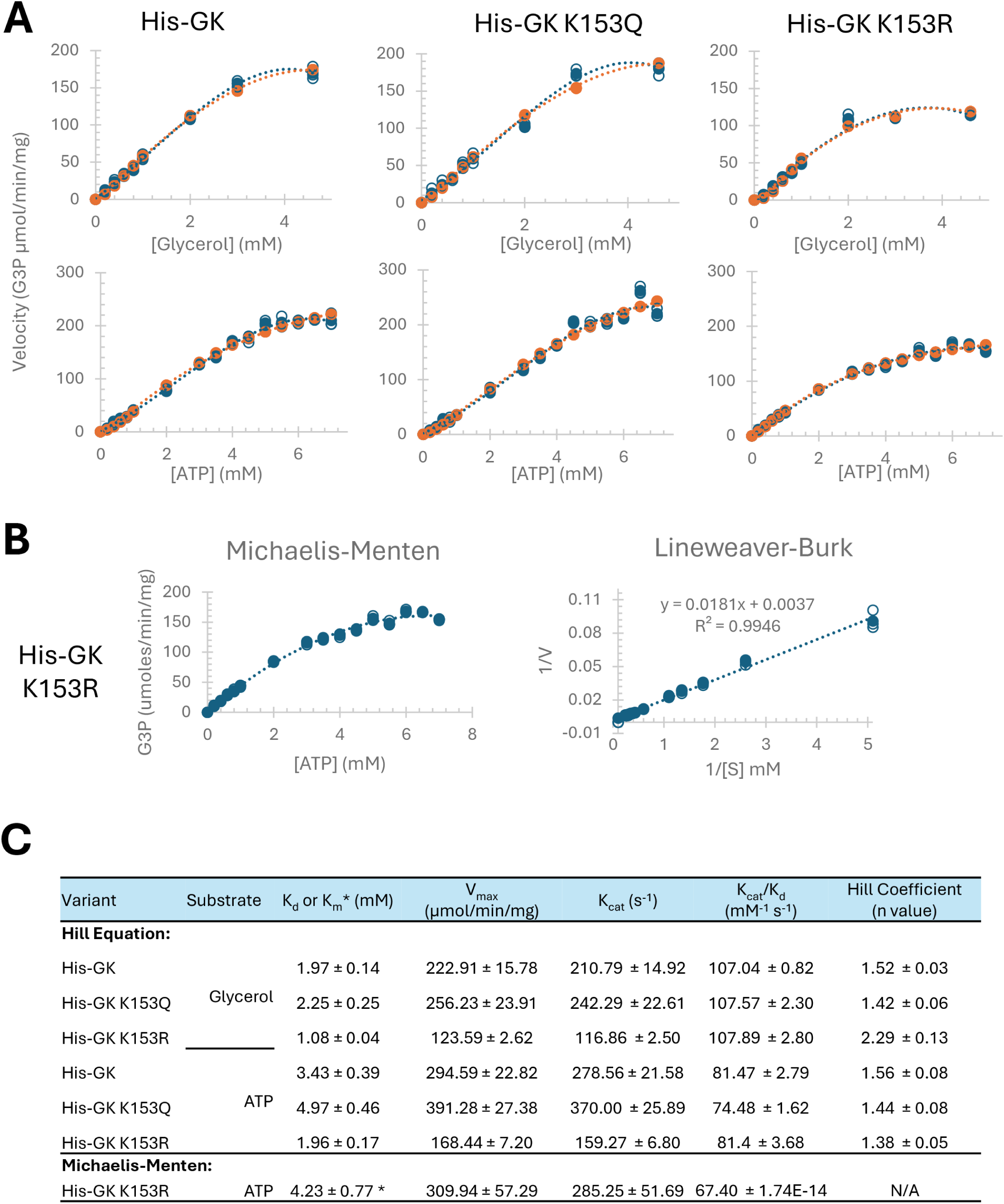
Kinetic analysis of His-GK variants purified from *H. volcanii* KM03 grown in GlyMM. A. Enzymatic activity of His-GK wt, K153Q (acetylation mimic), and K153R (non-acetylatable mutant) was measured under optimal assay conditions (50 mM HEPES, pH 8.0, 100 mM NaCl, 57 °C) using a coupled spectrophotometric assay that monitors NADH oxidation at 340 nm. Substrate concentrations were varied using glycerol and ATP. The kinetic profiles of His-GK wt and K153Q exhibited sigmoidal (non-Michaelis–Menten) behavior for both glycerol and ATP, consistent with cooperative substrate binding. In contrast, K153R showed Michaelis–Menten-like kinetics with ATP, indicating a loss of allosteric behavior. Experimental velocity data (blue) and best-fit model values (orange) are shown for each condition. B. Michaelis–Menten and Lineweaver–Burk plots for the K153R variant with ATP further support the non-cooperative kinetic behavior of this mutant. C. Summary of kinetic parameters: *K_m_, K_d_, V_max_*, *K_cat_*, *K_cat_*_/_*K_d_*, and Hill coefficient (*n*) for each enzyme-substrate combination. Data were fit using the Hill or Michaelis–Menten equations as appropriate. The Michaelis-Menten equation describes a simple enzyme-substrate interaction as v = (Vmax [S]) / (Km + [S]), where v is the reaction rate, Vmax is the maximum rate, [S] is the substrate concentration, and Km is the Michaelis constant. Hill equation used was Ycalculated = (Vmax)[L]n / (Kd + [L]n), where Y represents the fraction of occupied binding sites, [L] is the ligand concentration, n is the Hill coefficient indicating cooperativity, and Kd is the dissociation constant. Squared residuals (Yexperimental − Ycalculated)^2^ were summed to compute the sum of square residuals (SSR). Excel Solver was employed to minimize SSR and optimize the parameters for the best fit of the data. All measurements represent the mean of three biological replicates.

### Pat2-mediated acetylation of GK K153 appears to alter growth on glycerol

Our recent work demonstrates that Pat2 acetylates GK at K153 (23); however, the functional consequences of this modification remained unclear. To address this, we assessed the ability of His-GK or its variants to complement growth of a *ΔglpK* mutant in a Δ*pat2* background (KM06). Growth plate assays demonstrated that expression of either His-GK wt or the non-acetylated mimic K153R failed to rescue growth on GlyMM, whereas the acetylation mimic His-GK K153Q fully restored growth of the KM06 mutant under these conditions **(Fig 9A).** All constructs displayed growth on GluMM, confirming that the growth defect is specific to glycerol metabolism (**Fig 9B**). These findings highlight a previously unrecognized requirement for the K153 acetylation in enabling GK function under glycerol-utilizing conditions that is not required during growth on glucose.

**Fig 9.**
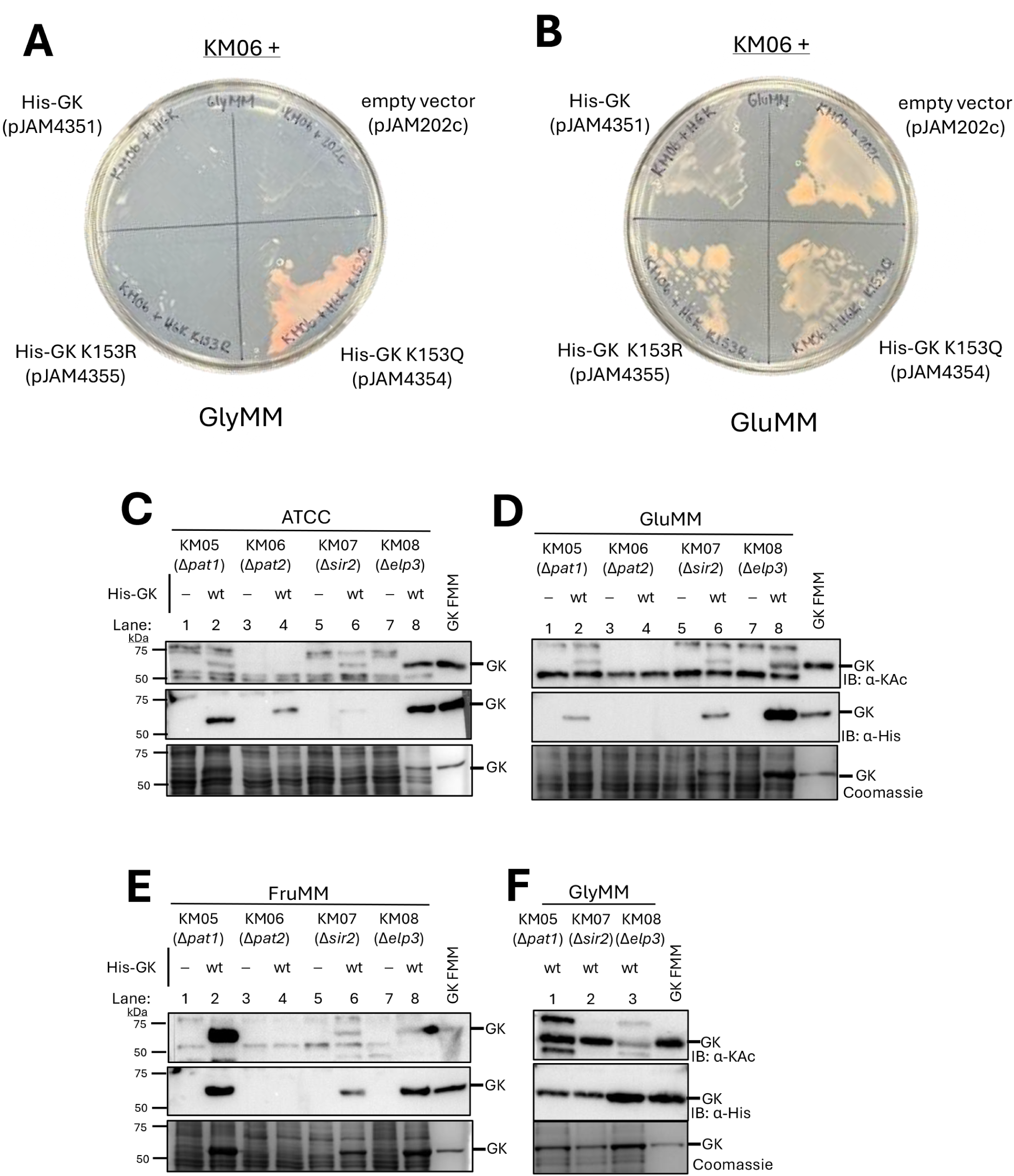
Pat2 acetylation or the K to Q acetylation mimic of the K153 site are associated with GK protein abundance and *H. volcanii* growth on glycerol. A. *Δpat2* Δ*glpK* mutant strains show growth defects on GlyMM when lacking GK (empty vector) or expressing His-GK wt or K153R from a plasmid, compared to the K153Q variant. KM06 (Δ*glpK* Δ*larC* Δ*pat2*) strains expressing His-GK wt, K153Q, K153R, or empty vector were streaked with loop onto GlyMM plates and incubated for 8 days. Growth was observed for the KM06 strain expressed the GK K153Q acetylation mimic compared to the other strains. B. Robust growth on GluMM observed for all *Δpat2 ΔglpK* mutant strains examined irrespective of the presence or absence of GK wt or variants. KM06 strains (as in panel A) were streaked onto on GluMM and incubated for 8 days. Growth was monitored by colony/streak formation. C-F. GK protein abundance and lysine acetylation occupancy are differentially affected by *Δpat2* and *Δsir2* mutations compared to *Δpat1* and *Δelp3*, with effects dependent on the carbon source. KM05 *(Δpat1*), KM06 *(Δpat2*), KM07 (*Δsir2*) and KM08 (*Δelp3*) strains expressing His-GK from plasmid pJAM4351 (wt) were compared to the empty vector control pJAM202c (–). The GNAT family (*pat1*, *pat2*, and *elp3*) and Sir2-type deacetylase (*sir2*) gene homolog mutations were generated in parent strain KM03 (H1207 *ΔglpK ΔlarC*). Cells were grown in different carbon sources as indicated: panels C (ATCC), D (GluMM), E (FruMM) and F (GlyMM). Whole cell lysate was normalized to 0.1 OD₆₀₀ per lane and lysed by boiling for 10 min in Laemmli SDS sample buffer (see Materials and Methods for details). Proteins were separated by SDS-PAGE. Equal protein loading was confirmed by CB stain. His-GK abundance and lysine acetylation were detected by IB: α-His and IB: α-KAc, respectively. His-GK purified from KM01 grown in FruMM was included as a positive control where indicated (GK FMM). KM06 and empty vector strains failed to grow in GlyMM, preventing protein analysis under these conditions (panel F). All experiments were performed in biological triplicates. Representative immunoblots are shown.

### Acetylation of GK K153 impacts GK abundance on glycerol

To further investigate the molecular basis of this phenotype, His-GK abundance and lysine acetylation status were determined in the presence of the GNAT-family acetyltransferase mutations Δ*pat1* (KM05), Δ*pat2* (KM06) and Δ*elp3* (KM08) and the Sir2-type deacetylase mutation Δ*sir2* (KM07). These strains were transformed with plasmids expressing His-GK wt or the empty vector. The K153Q and K153R variants were excluded from acetylation analysis, as these K153 mutations abolish the native acetylation site detected by immunoblot analysis. Cell pellets were analyzed by SDS-PAGE and immunoblotting **(Fig. 9C-F).** His-GK was detected and acetylated in KM05 (Δ*pat1*) across all carbon sources **(Fig. 9C–D** *lane 2* and **Fig. 9F** *lane 1*), with the highest acetylation levels observed in FruMM and GlyMM, indicating that *pat1* is not essential for GK acetylation. In KM06 (Δ*pat2*), His-GK acetylation was not detected and its protein abundance was barely visible in ATCC medium **(Fig. 9C**, *lane 4*) and undetectable in GluMM and FruMM **(Fig. 9D–F**, *lane 4*). Additionally, the strain failed to grow on GlyMM, suggesting that Pat2-mediated acetylation at K153 is positively correlated with GK protein abundance and that this impacts growth on glycerol. In contrast, KM07 (Δ*sir2*) and KM08 (Δ*elp3*) showed consistent GK acetylation regardless of the carbon source **(Fig. 9C–E**, *lanes 6 and 8*, **Fig. 9F** *lanes 2-3*), highlighting the specific and critical role of Pat2 in both GK acetylation and protein abundance.

## DISCUSSION

PTMs such as lysine acetylation have emerged as essential mechanisms regulating metabolic enzymes across all domains of life, including archaea (15). However, the functional significance of lysine acetylation in archaeal systems remains poorly understood. In this study, we characterized lysine acetylation of *H. volcanii* GK and identified K153 as the predominant acetylation site critical for enzyme function, abundance, stability, and carbon source adaptation.

### Dynamic acetylation of GK facilitates metabolic flexibility in *H. volcanii*

This study highlights the importance of GK acetylation dynamics in supporting the metabolic adaptability of *H. volcanii*. Carbon source availability strongly influenced the acetylated state of GK. Glycerol-grown cells exhibited a significantly higher abundance of acetylated GK compared to cells grown on glucose or rich (ATCC) media. A dynamic increase in acetylation occupancy at the K153 site was detected during the transition from glucose to glycerol. Functional analysis revealed that K153 acetylation was critical for metabolic adaptation: strains expressing His-GK or K153Q maintained growth following the carbon shift, whereas strains expressing the non-acetylated mimic K153R or empty vector experienced growth arrest.

These findings suggest that elevated GK acetylation under glycerol and fructose conditions reflects a metabolic prioritization strategy. In *H. volcanii*, glycerol and fructose promote rapid carbon flux toward pyruvate and acetyl-CoA, substrates that not only fuel metabolism but also enhance acetylation of key enzymes. In contrast, glucose and complex nutrient conditions may result in lower acetyl-CoA pools and reduced acetylation. This model aligns with previous studies showing that glycerol represses glucose metabolism via GlpR and favors carbon utilization strategies that enhance central flux and PTM-mediated regulation (7, 8, 28).

Dynamic shifts in protein acetylation regulating nutrient transitions are well documented in bacteria and eukaryotes. Lysine acetylation can alter the activities of key enzymes that balance carbon flux between glycolysis and gluconeogenesis and influence the branching between the citrate cycle and the glyoxylate bypass (29). Moreover, acetylation-driven changes can impact the levels of critical intermediary metabolites, including ATP, NAD^+^, and acetyl-CoA (30). These findings underscore that carbon source shifts require active and dynamic regulation rather than passive metabolic changes (31, 32). Acetylation is increasingly recognized as a global regulatory mechanism modulating metabolism, stress responses, and survival in prokaryotes (31, 33), further emphasizing the evolutionary conservation of PTMs in controlling key metabolic processes. Although PTMs in archaea are historically underappreciated (34), studies have shown that carbohydrate metabolism and glycerol utilization in archaea depend heavily on dynamic regulation (35, 36). Our AQUA-MS analysis demonstrated that K153 acetylation abundance correlates directly with enhanced GK enzymatic activity during glycerol growth. This functional coupling mirrors paradigms observed where nutrient availability drives PTM-mediated enzyme activation (29, 33). Collectively, these findings position acetylation at the K153 site of GK as a critical molecular switch for dynamic reprogramming of *H. volcanii* central metabolism during environmental nutrient shifts, further highlighting the ancient and conserved role of acetylation in regulating metabolic homeostasis.

Importantly, understanding the regulatory role of K153 acetylation extends beyond fundamental microbiology. Glycerol is a valuable compound in pharmaceutical, cosmetic, and food industries and is also a major byproduct of the biodiesel industry (37–39). Harnessing this surplus through metabolic engineering offers a sustainable route for bioconversion of this biodiesel industry byproduct into value-added compounds. Optimizing glycerol utilization through engineered enzymes, such as K153Q, which mimics the most active and stable acetylated form of GK, requires a detailed understanding of native regulatory mechanisms controlling key metabolic steps like glycerol phosphorylation (40, 41). These findings lay the groundwork for biotechnological applications that leverage archaeal enzymes in industrial biocatalysis and circular bioeconomy frameworks.

### Flexible loop-mediated regulation of GK via K153 acetylation

Here we find the primary site of GK acetylation, K153, is located within an apparent flexible loop, suggesting that GK activity may be regulated through loop-mediated structural dynamics. Flexible loops serve as pivotal regulatory elements in enzymes, modulating activity through conformational changes that can be controlled by PTMs. In *Homo sapiens*, phosphorylation of the conserved glycine-rich G-loop of protein kinases regulates ATP positioning and catalytic function in response to cellular signals (42). Furthermore, a recent study showed that acetylation of lysine residues within the flexible lid loop of the serine hydrolase ABHD17 significantly reduced its activity by restricting loop flexibility and substrate access, highlighting a direct structural mechanism of regulation (43). Like these examples, our study identifies a flexible loop that harbors K153 in *H. volcanii* GK that is conserved among most haloarchaea. We demonstrate that acetylation of K153 within this loop impacts enzymatic activity. This supports a model where PTMs targeting dynamic, conserved loops serve as a general mechanism for enzyme regulation across domains of life. The identification of a PTM-regulated loop in an archaeal enzyme extends loop-mediated regulatory concepts to extremophiles, suggesting that regulatory complexity via PTMs is more widespread in archaea than previously appreciated. Furthermore, understanding how flexible loops integrate metabolic signals through PTMs could inform future engineering of archaeal enzymes for biotechnological applications. Surface electrostatic modeling revealed that K153 resides within a neutral-to-positive region, suggesting accessibility for regulatory protein interactions, a pattern noted for other proteins (44). Consistent with broader trends, lysine acetylation predominates in membrane-interaction and dynamic loop regions (45) reinforcing the functional relevance of GK K153 acetylation.

### Evolutionary conservation of site-specific acetylation in enzyme regulation

While lysine acetylation sites are often difficult to predict based solely on amino acid sequence, owing to the diverse mechanisms that can lead to this PTM (14), the K153 site is highly conserved across diverse haloarchaeal GK homologs. While sequence alignment of GK homologs across the phylum *Euryarchaeota* show variable conservation of K153, this site is highly conserved within the class *Halobacteria*, indicating evolutionary specialization. Given that microorganisms must rapidly adjust metabolism to fluctuating carbon sources and that lysine acetylation can allow fast, reversible control of protein function (12, 46, 47), lysine acetylation at K153 may serve as a key adaptive mechanism in haloarchaea. Site-directed mutagenesis (K153Q and K153R) and *in vitro* acetylation assays reveal that K153 is the primary target of Pat2-mediated acetylation of GK, establishing a direct link between Pat2 and GK regulation. Similar mechanisms of site-specific acetylation controlling enzymatic activity have been documented in *Bacillus subtilis* AMP-forming acetyl-CoA synthetase (AcsA), where acetylation of a conserved lysine (K549) regulates nucleotide binding (48). In archaea, the chromatin protein Alba provides another well-characterized example: acetylation of a conserved lysine residue (K16) reduces its DNA-binding affinity, thereby modulating chromatin structure and gene expression in response to metabolic cues (49, 50). The conservation of lysine acetylation as a regulatory mechanism across bacteria, archaea, and eukaryotes suggests that modulation of enzyme activity via acetylation at structurally important lysine residues is an evolutionarily ancient strategy. Our discovery of K153-specific acetylation in *H. volcanii* GK thus expands the conserved paradigm of site-specific PTM regulation.

### Pat2-mediated acetylation stabilizes GK in *H. volcanii*

Our findings reveal that Pat2-mediated acetylation is essential for maintaining the stability of GK in *H. volcanii*. Deletion of *pat2* resulted in a complete loss of both GK protein abundance and acetylation signal, whereas the K153Q variant, which mimics constitutive acetylation, appeared of stable abundance even in the absence of Pat2. Furthermore, thermal shift analysis revealed that acetylation at K153 enhances the thermal stability of *H. volcanii* GK, particularly in the presence of its physiological ligands. The acetylation mimic K153Q consistently exhibited higher Tm values than the non-acetylated mimic K153R across the tested conditions, indicating greater structural stability. These results suggest that acetylation at K153, mediated by Pat2, serves not only as an enzymatic activity regulator, but may also play a stabilizing and protective role in preventing GK degradation.

This mechanism, in which lysine acetylation appears to stabilize the haloarchaeal GK, parallels regulatory strategies observed in other systems. In *Homo sapiens*, the acetyltransferase p300 stabilizes homeodomain-interacting protein kinase 2 (HIPK2) by promoting acetylation, which prevents ubiquitin-mediated proteasomal degradation and amplifies tumor suppressor function (51). Similarly, acetylation at K751 of large tumor suppressor kinase 1 (LATS1) stabilizes the protein by blocking ubiquitination, although it simultaneously inhibits kinase activation, shifting LATS1 function as a tumor suppressor to an oncoprotein (52). Glycogen synthase kinase-3β (GSK-3β) stability is also regulated by tau-induced acetylation at K15, which blocks ubiquitin-mediated proteolysis and leads to elevated kinase activity, contributing to a pathogenic feedback loop in Alzheimer’s disease (53).

Collectively, these examples highlight a conserved regulatory principle across domains of life: acetylation serves as a critical mechanism to stabilize key metabolic or protein kinases by protecting them from degradation. In *H. volcanii*, Pat2-mediated acetylation of GK similarly ensures appropriate GK levels to support metabolic flexibility during environmental shifts. These findings expand the emerging understanding of post-translational regulation in archaea and suggest that acetylation-mediated stabilization mechanisms may be broadly conserved, even among extremophilic organisms.

### Acetylation-Dependent Oligomeric Transitions in GK

PTMs such as lysine acetylation have been increasingly recognized as regulators of enzyme oligomerization and function. In *Helicobacter pylori*, acetylation modulates the oligomeric state of RNase J, promoting its assembly into higher-order complexes that are critical for ribonucleolytic activity (54). Widespread acetylation of metabolic enzymes has also been documented across diverse organisms, including both cytosolic and mitochondrial proteins (55). However, the functional consequences of acetylation are often enzyme specific.

In this study, we demonstrate that acetylation at lysine 153 plays a critical role in regulating the activity, allosteric behavior, and oligomeric state of GK from *H. volcanii*. SEC analysis revealed that all GK variants remain dimeric under non-glycerol conditions (glucose, fructose, and complex media). However, under high-acetylation conditions—specifically when cells are grown on glycerol as the sole carbon source—wild-type GK also forms a tetrameric species (3), suggesting an acetylation-dependent oligomeric transition.

Consistent with this model, our kinetic analyses revealed that only the wild-type and K153Q (acetylation mimic) variants retained sigmoidal behavior with both glycerol and ATP substrates, reflecting the positive cooperativity previously reported for *H. volcanii* His-GK. In contrast, the non-acetylatable K153R variant exhibited reduced activity across both substrates and a clear shift toward Michaelis–Menten kinetics with ATP. This transition from cooperative to non-cooperative behavior underscores the functional significance of acetylation at K153 in maintaining an allosterically responsive conformation, particularly for ATP binding.

The elevated Hill coefficient observed for K153R with glycerol likely reflects a distorted or compensatory conformational response, rather than enhanced allosteric regulation. This aberrant cooperativity, coupled with the ATP-specific loss of sigmoidal kinetics, points to allosteric misregulation resulting from impaired acetylation at K153. These findings align with our thermal shift assay data, which showed that the K153R variant has significantly reduced thermal stability in the presence of ATP, indicative of altered conformational flexibility or structural instability.

Taken together, our data support a model in which K153 acetylation functions as a molecular switch, enabling GK to transition into an allosterically competent, tetrameric state that responds dynamically to substrate levels. In the absence of acetylation (K153R), the enzyme retains basic catalytic activity but loses the cooperative behavior characteristic of a finely tuned metabolic sensor. This change reflects a broader architectural disruption, linking acetylation to both conformational plasticity and oligomeric state.

These results highlight K153 acetylation as a critical regulatory point in GK function, bridging post-translational modification with enzyme dynamics. This mechanism may serve as a metabolic tuning strategy in *H. volcanii*, allowing the organism to optimize glycerol metabolism under fluctuating environmental conditions through modulation of enzyme conformation, cooperativity, and activity.

## CONCLUSION

This study reveals that lysine acetylation at K153 is essential for *H. volcanii* GK function, stability, and metabolic adaptation. Acetylation levels dynamically increased during glycerol utilization, correlating with enhanced GK enzymatic activity, while mutations abolishing acetylation impaired enzyme function and hindered growth under carbon source transition conditions. Furthermore, our kinetic analyses further demonstrated that acetylation at K153 is required to maintain allosteric behavior, particularly with ATP, linking this post-translational modification to the enzyme’s ability to sense and respond to substrate levels. Our findings expand the current understanding of post-translational regulation in archaea and demonstrate that lysine acetylation serves as a key mechanism for tuning enzyme activity, stability, and structural dynamics in response to environmental fluctuations. These results establish acetylation as a critical regulatory strategy for metabolic flexibility in *H. volcanii* and highlight the broader significance of PTMs in archaeal adaptation. Future work investigating how acetylation integrates with other regulatory networks will further elucidate the complexity of metabolic control in extremophilic archaea and inform potential biotechnological applications leveraging archaeal enzyme stability and regulation.

## Acknowledgments

Funds awarded to JMF to characterize archaeal biocatalysts were through the U.S. Department of Energy, Office of Basic Energy Sciences, Division of Chemical Sciences, Geosciences and Biosciences, Physical Biosciences Program (DOE DE-FG02-05ER15650) and to determine fundamental biological mechanisms related to human health were through National Institutes of Health (NIH R01 GM57498). Proteomics experiments were supported by the Mass Spectrometry Research and Education Center at the University of Florida, funded by NIH grants S10 OD021758-01A1 and S10 OD030250-01A1. The PRM mass spec analysis was supported by the U.S. National Science Foundation grant number 2414925 (Original award number 2042182) and the start-up funding from the University of Florida to XW.

